# Developmental change in prefrontal cortex recruitment supports the emergence of value-guided memory

**DOI:** 10.1101/2021.02.13.431073

**Authors:** Kate Nussenbaum, Catherine A. Hartley

## Abstract

Prioritizing memory for valuable information can promote adaptive behavior across the lifespan, but it is unclear how the neurocognitive mechanisms that enable the selective acquisition of useful knowledge develop. Here, using a novel task coupled with functional magnetic resonance imaging, we examined how children, adolescents, and adults (N = 90) learn from experience what information is likely to be rewarding, and modulate encoding and retrieval processes accordingly. We found that the ability to use learned value signals to selectively enhance memory for useful information strengthened throughout childhood and into adolescence. Encoding and retrieval of high- vs. low-value information was associated with increased activation in striatal and prefrontal regions implicated in value processing and cognitive control. Age-related increases in value-based lateral prefrontal cortex modulation mediated the relation between age and memory selectivity. Our findings demonstrate that developmental increases in the strategic engagement of the prefrontal cortex support the emergence of adaptive memory.

## Introduction

Memories of past experiences guide our behavior, promoting adaptive action selection throughout our lives (Biderman et al., 2020). But not all experiences are equally useful to remember — the information we encounter varies in its utility in helping us gain future reward. By adulthood, individuals demonstrate the ability to prioritize memory for information that is likely to be most rewarding in the future (Adcock et al., 2006; M. S. Cohen et al., 2019, 2014; Hennessee et al., 2019; Shigemune et al., 2014; Shohamy & Adcock, 2010; Wittmann et al., 2005). Children, however, demonstrate weaker memory selectivity, often remembering relatively inconsequential information at the expense of higher-value items or associations (Castel et al., 2011; Hanten et al., 2007; Nussenbaum et al., 2020). Behavioral studies have found that the use of value to guide encoding and retrieval processes emerges and strengthens gradually throughout childhood and adolescence, promoting more efficient acquisition of useful knowledge with increasing age (Castel et al., 2011; Hanten et al., 2007; Nussenbaum et al., 2020). It is unclear, however, how changes in brain activity support this observed emergence of motivated memory. Though a large literature has examined developmental change in the neural mechanisms that support memory from early childhood to young adulthood (Ghetti & Fandakova, 2020; Ofen, 2012; Shing et al., 2010), no prior studies have investigated how the developing brain prioritizes memories based on their relative utility.

Prioritizing valuable information in memory requires both determining the value of information and strategically modulating encoding accordingly. The vast majority of adult studies have focused only on the strategic *use* of value to guide memory — in most studies of motivated memory, *determining* the value of information is trivial for participants because experimenters label to-be-remembered information with explicit value cues (e.g., dollar signs, stars, point amounts) (Adcock et al., 2006; Castel et al., 2011; M. S. Cohen et al., 2014; Murty et al., 2017). However, in real-world contexts, individuals must derive the value of information from the statistics of their environments. In a recent behavioral study (Nussenbaum et al., 2020), we demonstrated that young adults could use naturalistic value signals to prioritize memory for useful information. Specifically, we manipulated information value via item *frequency*. Across many environments, the frequency of encountering something in the past predicts the frequency of encountering it in the future. In this way, frequency can signal information utility (Anderson & Milson, 1989; Anderson & Schooler, 1991; Liu et al., 2020; Pachur et al., 2014; Rich & Gureckis, 2018; Stevens et al., 2016) — if individuals are likely to encounter something often in the future, encoding information about it is likely to be valuable. For example, remembering the supermarket aisle of an ingredient with which one often cooks is likely to facilitate greater reward gain than remembering the aisle of an ingredient one almost never uses. In our prior study, we translated this feature of real-world environments to a laboratory task in which individuals could learn the potential reward value of associative information by first learning the relative frequency of items in their environments. We found that individuals exploited these naturalistic value signals and demonstrated better memory for information associated with high-relative to low-frequency items. Critically, this pattern of results varied with age; the strategic prioritization of high-value information in memory increased from age 7 to age 25 (Nussenbaum et al., 2020). It is unclear, however, if developmental improvements in memory prioritization stemmed from differences in learning the relative value of information based on environmental statistics or *using* learned value signals to strategically prioritize memory. Each of these processes likely engages separable neural systems.

Deriving value from the structure of the environment first requires the learning of statistical regularities. In the case of learning the frequency with which one might need to use information, individuals must differentiate novel occurrences (e.g., cooking with a rare ingredient) from oft-repeated experiences (e.g., cooking with a common food). Neurally, medial temporal lobe regions may support sensitivity to item repetitions. The parahippocampal cortex in particular demonstrates reduced responsivity to repeated relative to novel presentations of items (i.e., “repetition suppression”) (Gonsalves et al., 2005; Kirchhoff et al., 2000; Köhler et al., 2005; O’Kane et al., 2005; Turk-Browne et al., 2006). Though some accounts of repetition suppression suggest that attenuated responses simply indicate neural “fatigue,” the phenomenon has also been shown to be sensitive to the statistical context of the environment, suggesting that suppression may reflect stimulus expectation and index learning of environmental regularities (Auksztulewicz & Friston, 2016). Repetition suppression has also been shown to relate to implicit memory for repeated items (Ward et al., 2013). Paralleling their robust implicit learning abilities (Amso & Davidow, 2012; Finn et al., 2016; Meulemans et al., 1998), children and adolescents also demonstrate neural repetition suppression effects (Nordt et al., 2016; Scherf et al., 2011; Turi et al., 2015), though repetition suppression — and the ability to learn the statistical structure of the environment — may increase throughout childhood (Scherf et al., 2011). When individuals need to remember information associated with previously encountered stimuli (e.g., the grocery store aisle where an ingredient is located), frequency knowledge may be instantiated as value signals, engaging regions along the mesolimbic dopamine pathway that have been implicated in reward anticipation and the encoding of stimulus and action values. These areas include the ventral tegmental area (VTA) and the ventral and dorsal striatum (Adcock et al., 2006; Liljeholm & O’Doherty, 2012; Shigemune et al., 2014).

*Using* these learned value signals to guide memory likely requires cognitive control (Castel et al., 2007; M. S. Cohen et al., 2014). Value responses in the striatum may signal the need for increased engagement of the dorsolateral prefrontal cortex (dlPFC) (Botvinick & Braver, 2015), which supports the implementation of strategic control. Enhanced recruitment of control processes promotes the use of deeper and more elaborative encoding strategies (M. S. Cohen et al., 2019, 2014; Miotto et al., 2006; Uncapher & Wagner, 2009) as well as the selection and maintenance of effective retrieval and post-retrieval monitoring strategies (Libby & Lipe, 1992; Scimeca & Badre, 2012), which may contribute to better memory for high-value information. The use of value to proactively upregulate cognitive control responses improves throughout development, though the specific trajectory of improvement may relate to the control demands of a given task (Davidow et al., 2018). Selectively enhancing the use of encoding and retrieval strategies requires not only tight coordination between subcortical regions involved in value processing and prefrontal areas implicated in control (Murty & Adcock, 2014), but also an available repertoire of memory strategies to implement. Even in the absence of value cues, children and adolescents demonstrate reduced use of strategic control (Bjorkland et al., 2009) and reduced lateral prefrontal engagement during encoding (Ghetti et al., 2010; Ghetti & Fandakova, 2020; Shing et al., 2016; Tang et al., 2018), suggesting that the availability of mnemonic control strategies may increase with age.

Taken together, prior work suggests that adaptive memory requires the recruitment and coordination of multiple neural systems, including mechanisms for learning environmental structure, representing value, and engaging strategic control, all of which may undergo marked changes from childhood to adulthood. Here, we examined how the development of these neurocognitive processes supports the emergence and strengthening of value-guided memory from childhood to young adulthood. To pinpoint loci of developmental differences in adaptive memory prioritization, we combined our novel motivated memory experiment (Nussenbaum et al., 2020) with functional neuroimaging. During the task, participants first learned the frequency of items in their environments, and then learned information associated with each item. Importantly, we structured our task such that the frequency with which participants first experienced each item indicated the frequency with which they would be asked to report the information associated with it, and therefore, the number of points they could earn by remembering the association. Immediately following encoding, we administered a memory test in which participants had to select each item’s correct associate. Because frequency of exposure to an item may facilitate subsequent associative memory even when it does *not* signal the value of information (Popov & Reder, 2020; Reder et al., 2016), in our prior behavioral study (Nussenbaum et al., 2020), we examined the effects of item frequency on subsequent associative memory in two contexts: one in which item frequency signaled information value and one in which it did not. Critically, we found that with our experimental design, frequency only facilitated memory when it signaled the value of remembering information — increased item exposure did not in and of itself enhance subsequent associative memory. Thus, in the present fMRI study, we focused only on the condition in which item frequency *did* indicate the potential reward that could be earned for remembering associations.

We examined neural activation during the learning of item frequency, and when participants were asked to encode and retrieve information associated with high- vs. low-frequency items. We hypothesized that while participants across our entire age range would demonstrate sensitivity to the frequency of items in their environments, with increasing age, participants would show improvements in transforming this experiential learning into value signals and modulating the engagement of strategic control processes during encoding. Neurally, we expected that at encoding and retrieval, the recognition of information value would be reflected in increased striatal activation in response to associations involving high- vs. low-frequency items, while the engagement of strategic control would be reflected in increased activation in lateral prefrontal cortex. Further, we hypothesized that increased recruitment of the striatum and prefrontal cortex during encoding and retrieval of high- vs. low-value information would underpin the strengthening of adaptive memory prioritization from childhood to early adulthood.

## Results

### Approach

Participants ages 8 – 25 years (N = 90; 30 children ages 8 – 12 years; 30 adolescents ages 13 – 17 years; 30 adults ages 18 – 25 years) completed two blocks of three tasks (Figure 1) while undergoing functional magnetic resonance imaging. In the first, *frequency-learning* task, participants viewed a continuous stream of 24 unique postcards, one at a time. Twelve of the postcards only appeared once, while 12 repeated five times. Participants indicated whether each postcard they viewed was old or new. In the second, *associative encoding* task, participants viewed the type of stamp that went on each type of postcard. Participants were instructed that in the subsequent task, they would have to stamp *all* of their postcards, earning one point for each postcard stamped correctly. Critically, in the associative encoding task, regardless of the number of each type of postcard that they had (i.e., 1 or 5), participants saw each type of postcard with its corresponding stamp only *once*. Thus participants were informed that the prior frequency of each postcard indicated the value of encoding its associated stamp, but they had equal exposure to the to-be-encoded associations across frequency conditions. In the *retrieval* task, participants had to indicate the stamp that went with each unique postcard from one of four options, earning one point for each postcard stamped correctly. After stamping each unique postcard once, participants were asked to report its original frequency on a scale from 1 to 7. Finally, participants stamped all remaining postcards, such that they completed 48 additional memory test trials (i.e. they stamped each of the postcards in the 5-frequency condition four more times.) These trials were not included in any analyses, but their inclusion ensured that correctly encoding the stamps that belonged on the high-frequency postcards would be more valuable for participants despite each retrieval trial being worth one point. After completing the three tasks, participants were told that they were going to play a second set of similar games. The second set of tasks was identical to the first, except that the stimuli were changed from postcards and stamps to landscape pictures and picture frames. The order of the stimulus sets was counterbalanced across participants, and all data were combined across blocks.

**Figure 1.**
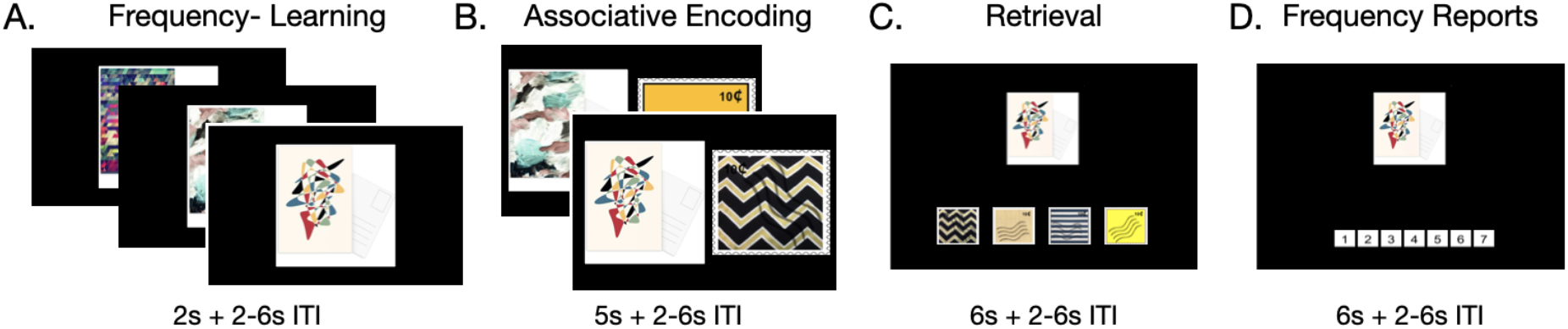
Task structure. Participants first learned the frequencies of each item (A) by viewing them in a continuous stream. They then were shown the information associated with each item (B). During retrieval, participants had to report the information associated with each item (C) as well as the item’s original frequency (D).

Across our behavioral analyses, we treated age as a continuous variable. To test for nonlinear effects of age, we first compared the fit of models with a linear age term and with both a linear and quadratic age term (Braams et al., 2015; Somerville et al., 2013). We dropped the quadratic age term when it did not significantly improve model fit. Because this study was cross-sectional, one concern was that the children, adolescents, and adults that we recruited may have come from different populations. Indeed, we observed a significant relation between age and age-normed Wechsler Abbreviated Scale of Intelligence (WASI; (Wechsler, 2011) scores in our sample (β = −.60, SE = .26, *p* = .0238), suggesting the children had slightly higher estimated IQs for their age relative to adults. To account for these age-related differences in reasoning ability, we included age-normed WASI scores as an interacting fixed effect in all analyses. Our aim in including WASI scores as a control variable was to partially account for confounding, population-level differences across our age groups, enabling us to more clearly examine the relation between age itself and our neurocognitive processes of interest.

### Experiential learning of environmental statistics improved with age

During frequency learning, participants across our age range responded to new and repeated items with a high degree of accuracy (new items: *mean* = .90, *SD* = .30; repeated items: *mean* = .92, *SD* = .27; Figure S2A). Older participants demonstrated higher accuracy in correctly identifying both new and repeated items (generalized mixed-effects model results: *new items:* X^2^(1) = 25.52, *p* <.001, *repeated items:* X^2^(1) = 33.43, *p* < .001). Participants were also more accurate in identifying items as ‘repeated’ as the number of times they saw each item increased, *χ*^2^ = 138.03, *p* < .001, indicating learning throughout the task. This effect varied as a function of age — younger participants demonstrated a larger effect of the number of item repetitions on response accuracy, as indicated by a significant interaction, χ^2^(1) = 17.41, *p* < .001.

Response times to both new and old items decreased with age (Figure S2B; linear mixed-effects model results*: new items: F*(1, 85.99) = 32.51, *p* < .001; *old items:* F(1, 87.55) = 21.82, *p* < .001), such that reaction times decreased steeply throughout childhood before leveling off into late adolescence and early adulthood. Finally, response times for old items also decreased as the number of item repetitions increased, *F*(1, 69.94) = 282.21, *p* < .001.

Participants’ ability to distinguish old from new items was associated with a wide network of neural regions, some of which demonstrated greater activation in response to the last vs. first appearance of each item, and others of which demonstrated suppressed activation across repetitions. Specifically, whole-brain contrasts revealed greater recruitment of regions of the lateral occipital cortex, the frontal pole, precuneus, angular gyrus, and caudate (among other regions, see Figure 2A and Supplementary Table S29), on the last vs. first appearance of each item. We observed widespread repetition suppression effects, reflected in *decreases* in neural responsivity in the lateral occipital cortex and temporal occipital cortex on the last vs. first appearance of each item. In line with our hypothesis, we also observed a robust decrease in activation in the parahippocampal cortex (Figure 2B, Supplementary Table S29).

**Figure 2:**
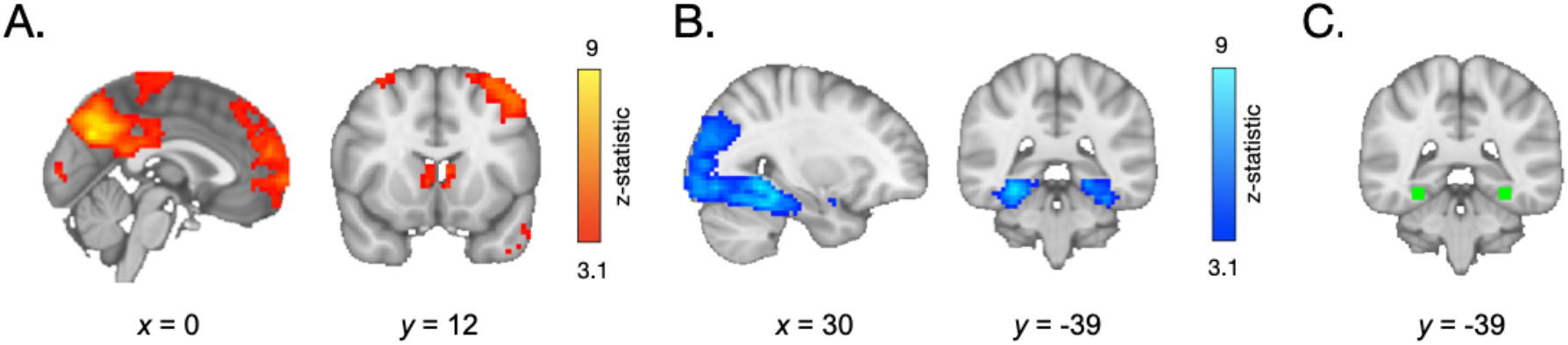
(A) During frequency-learning, participants demonstrated increased recruitment of regions in the frontal cortex, angular gyrus, and striatum on the last vs. first appearance of high-frequency items. (B) They demonstrated *decreased* activation in the lateral occipital cortex, temporal occipital cortex, and parahippocampal cortex. (C) Within a parahippocampal ROI (shown in green), the decrease in responses to each stimulus on its last vs. first appearance was greater in older participants.

We next examined whether repetition suppression in the parahippocampal cortex changed with age. We defined a parahippocampal region of interest (ROI) by drawing a 5mm sphere around the peak voxel from the group-level first > last appearance contrast (*x* = 30, *y* = −39, *z* = −15), and mirrored it to encompass both right and left parahippocampal cortex (Figure 2C). For each participant, we modeled the neural response to each appearance of each high-frequency item. We then examined how neural activation changed as a function of repetition number and age. To account for nonlinear effects of repetition number, we included linear and quadratic repetition number terms. In line with our whole-brain analysis, we observed a main effect of repetition number, *F*(1, 5015.9) = 30.64, *p* < .001, indicating that neural activation within the parahippocampal ROI decreased across repetitions. Further, we observed a main effect of quadratic repetition number, *F*(1, 9881.0) = 7.47, *p* = .006, indicating that the reduction in neural activity was greatest across earlier repetitions (Fig 3A). Importantly, the influence of repetition number on neural activation varied with both linear age, *F*(1, 7267.5) = 7.2, *p* = .007, and quadratic age, *F*(1, 7260.8) = 6.9, *p* = .009. Finally, we also observed interactions between quadratic repetition number and both linear and quadratic age (*p*s < .026). These age-related differences suggest that repetition suppression was greatest in adulthood, with the steepest increases occurring from late adolescence to early adulthood (Figure 3).

**Fig 3.**
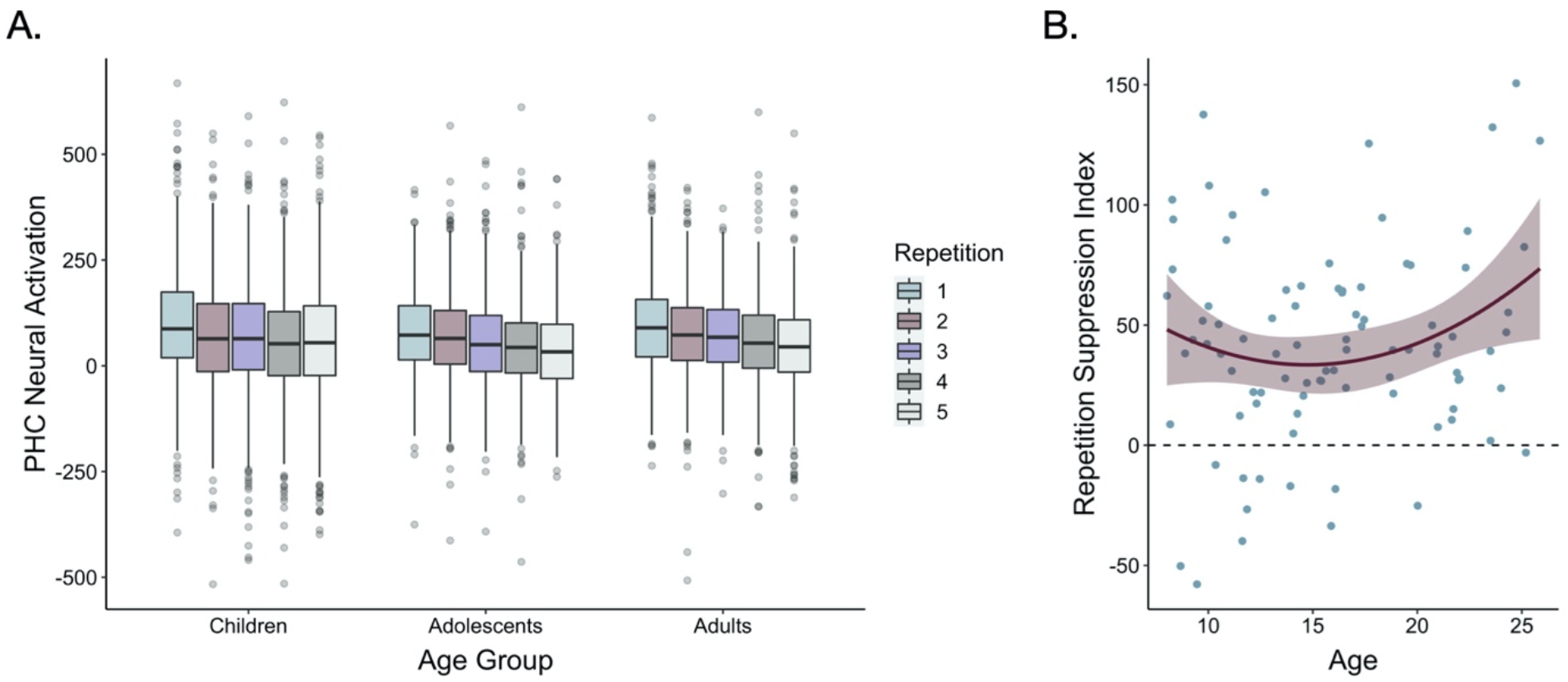
(A) Neural activation within a bilateral parahippocampal cortex ROI decreased across stimulus repetitions both linearly, *F*(1, 5015.9) = 30.64, *p* < .001, and quadratically, *F*(1, 9881.0) = 7.47, *p* = .006. Repetition suppression increased with linear age, *F*(1, 7267.5) = 7.2, *p* = .007, and quadratic age *F*(1, 7260.8) = 6.9, *p* = .009. The horizontal black lines indicate median neural activation values. The lower and upper edges of the boxes indicate the first and third quartiles of the grouped data, and the vertical lines extend to the smallest value no further than 1.5 times the interquartile range. Grey dots indicate data points outside those values. (B) The decrease in neural activation in the bilateral PHC ROI from the first to fifth repetition of each item also increased with both linear age, *F*(1, 78.32) = 3.97, *p* = .05, and quadratic age, *F*(1, 77.55) = 4.8, *p* = .031.

For each participant for each item, we also computed a “repetition suppression index” by taking the difference in mean beta values within our ROI on each item’s first and last appearance (Ward et al., 2013). These indices demonstrated a similar pattern of age-related variance — we found that the reduction of neural activity from the first to last appearance of the items varied positively with linear age, *F*(1, 78.32) = 3.97, *p* = .05, and negatively with quadratic age, *F*(1, 77.55) = 4.8, *p* = .031 (Figure 3B). Taken together, our behavioral and neural results suggest that sensitivity to the repetition of items in the environment was prevalent from childhood to adulthood but increased with age.

### Age-related differences in explicit knowledge of environmental structure

Could participants transform their sensitivity to environmental statistics into explicit reports of item frequency? To address this question, we computed participants’ frequency report error magnitudes by taking the absolute value of the difference between the item’s true frequency (i.e., 1 or 5) and each participant’s explicit report of its frequency (i.e., 1 - 7). We then examined how these report error magnitudes varied as a function of age and frequency condition. We observed a main effect of age (*F*(1, 94.30) = 17.57, *p* < .001) such that error magnitudes decreased with increasing age (Children: *Mean* = 1.48, *SD* = 1.34; Adolescents: *Mean* = 1.10, *SD* = 1.12; Adults: *Mean* = 1.13, *SD* = 1.05). Error magnitudes were not related to frequency condition (*p* = .993), indicating that participants were not systematically better at representing the ‘true’ frequencies of items that appeared once or items that appeared five times.

To examine relations between online frequency learning and explicit knowledge, we tested whether repetition suppression indices for each item related to frequency reports. We hypothesized that participants would report the items that elicited the greatest repetition suppression as the most frequent. However, in line with other studies suggesting dissociations between repetition suppression and explicit memory (Ward et al., 2013), we did not observe any relation between repetition suppression indices and frequency reports, *F*(1, 1360.74) = 0.01, *p* = .903. Thus, while we observed parallel developmental improvements in online frequency learning and subsequent explicit reports, they may be driven by separable processes.

### Age-related differences in value-guided memory

Participants’ frequency-learning performance and their explicit frequency reports indicate that older participants were better both at tracking repetitions of items within their environments and at explicitly representing item frequencies. Were participants able to use these representations of the structure of their environment to prioritize memory for high-value information? To address this question, we examined how frequency condition and age influenced memory accuracy. Memory accuracy varied as a function of both linear (χ^2^(1) = 8.68, *p* = .003) and quadratic age (χ^2^(1) = 4.24, *p* = .039), such that older participants demonstrated higher memory accuracy, with the steepest improvements in memory accuracy occurring from childhood into early adolescence (Figure 4). In line with our hypothesis, we observed a main effect of frequency condition on memory, χ^2^(1) = 19.73, *p* <.001, indicating that individuals used naturalistic value signals to prioritize memory for high-value information. Critically, this effect interacted with both linear age (χ^2^(1) = 10.74, *p* = .001) and quadratic age (χ^2^(1) = 9.27, *p* = .002), such that the influence of frequency condition on memory increased to the greatest extent throughout childhood and early adolescence.

**Figure 4.**
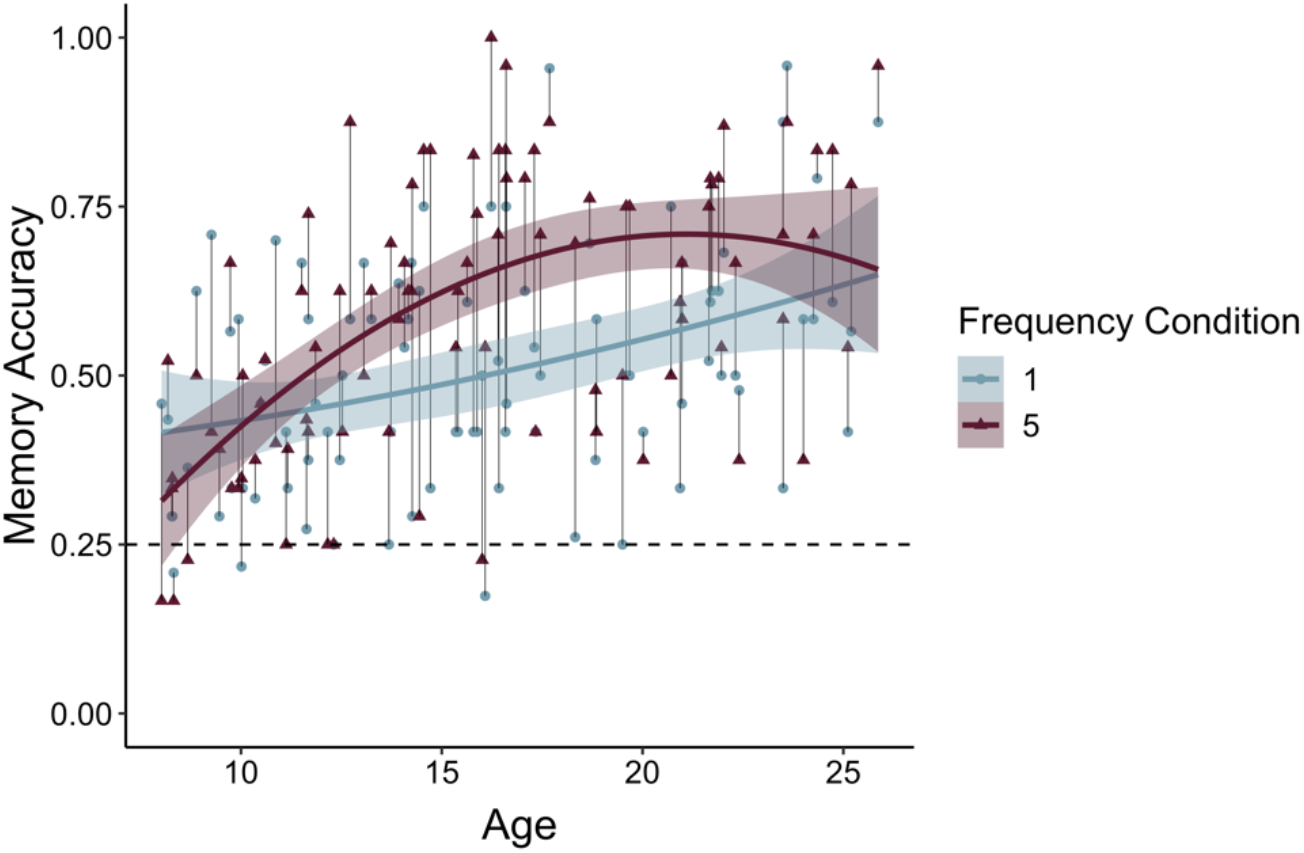
Participants demonstrated prioritization of memory for high-value information, as indicated by higher memory accuracy for associations involving items in the five-relative to the one-frequency condition (χ^2^(1) = 19.73, *p* <.001). The effects of item frequency on associative memory increased throughout childhood and into adolescence (linear age x frequency condition: χ^2^(1) = 10.74, *p* = .001; quadratic age x frequency condition: χ^2^(1) = 9.27, *p* = .002).

To determine whether the interaction between quadratic age and frequency condition on memory accuracy reflected an adolescent peak in value-guided memory prioritization, we re-ran our memory accuracy model without including any age terms and extracted each participant’s random slope across frequency conditions. We then submitted these random slopes to the “two-lines” test (Simonsohn, 2018), which fits two regression lines with oppositely signed slopes to the data, algorithmically determining where the sign flip should occur. The results of this analysis revealed that the influence of frequency condition on memory significantly increased from age 8 to age 15.86 (b = .03, *z* = 2.71, *p* = .0068; supplementary Figure S3), but only marginally decreased from age 15.86 to age 25 (b = −.02, *z* = 1.91, *p* = .0576). Thus, the interaction between frequency condition and quadratic age on memory performance suggests that the biggest age differences in value-guided memory occurred through childhood and early adolescence, with older adolescents and adults performing similarly.

Because we observed age-related differences in participants’ online learning of item frequencies and in their explicit frequency reports, we further examined whether these age differences in initial learning could account for the age differences we observed in associative memory. To do so, we ran an additional model in which we included each participant’s mean frequency learning accuracy, mean frequency learning accuracy on the last repetition of each item, and explicit report error magnitude as covariates. Here, explicit report error magnitude predicted overall memory performance, X^2^(1) =13.05, *p* < .001, and we did not observe main effects of age or quadratic age on memory performance (*p*s > .20). However, we continued to observe a main effect of frequency condition, X^2^(1) = 19.65 *p* < .001, as well as significant interactions between frequency condition and both linear age X^2^(1) = 10.59, *p* = .001, and quadratic age X^2^(1) = 9.15, *p* = .002. Thus, while age differences in initial learning related to overall memory performance, they did not account for age differences in the use of environmental regularities to strategically prioritize memory for valuable information.

### Neural mechanisms of value-guided encoding

We next examined how neural activation during encoding supported the use of learned value to guide memory. Specifically, we examined whether participants demonstrated different patterns of neural activation during encoding of information associated with high- vs. low-frequency items. In line with our hypothesis, a whole-brain contrast revealed increased engagement of the left lateral PFC and bilateral caudate (1765 voxels at *x* = −51, *y* = 42, *z* = 9; 232 voxels at *x* = 18, *y* = 12, *z* = 6; and 54 voxels at *x* = 18, *y* = 18, and *z* = 12; Figure 5A) during encoding of the pairs involving high-frequency items relative to pairs involving low-frequency items. To examine how this pattern of activation related to behavior, we computed a ‘memory difference score’ for each participant by subtracting their memory accuracy for associations involving low-frequency items from their accuracy for associations involving high-frequency items. We then included these memory difference scores as a covariate in our group-level GLM examining neural activation during encoding of pairs involving high- vs. low-frequency items. Participants who demonstrated the greatest difference in memory accuracy for pairs involving high-frequency vs. low-frequency items also demonstrated greater value-based modulation of left lateral PFC activation (232 voxels at *x* = −48, *y* = 21, *z* = 27; Figure 5B).

**Figure 5:**
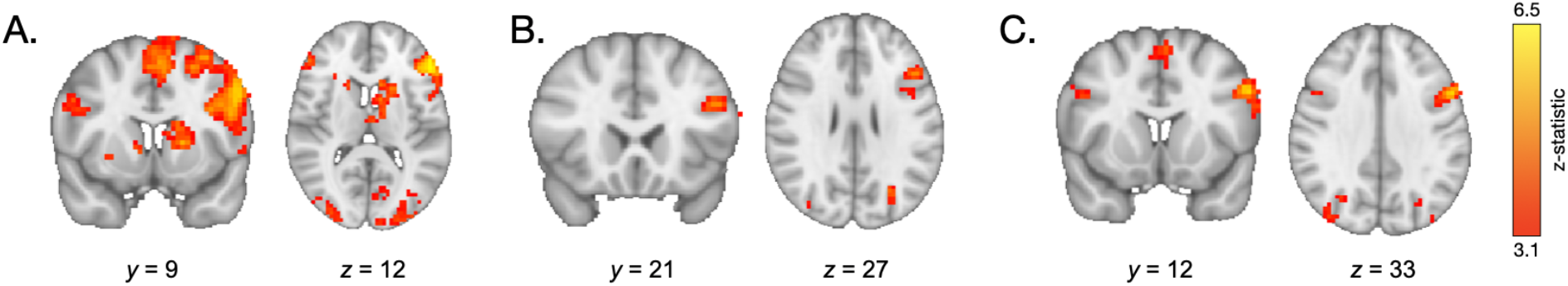
(A) During encoding of associations involving high- vs. low-frequency items, participants demonstrated greater engagement of the lateral PFC and caudate. (B) Participants who demonstrated the greatest value-based modulation of memory also demonstrated the greatest modulation of left prefrontal cortical activation during encoding of high- vs. low-value associations. (C) During encoding of both high- and low-value pairs, older participants demonstrated greater recruitment of the PFC relative to younger participants.

Because participants demonstrated effects of value on memory, neural signatures of encoding high- vs. low-value information may reflect successful vs. unsuccessful encoding. To de-confound the effects of value vs. subsequent memory accuracy on neural activation at encoding, we re-ran our high- vs. low-value contrast but restricted our analysis to associations that were subsequently retrieved correctly. We observed similar neural effects — increased recruitment of the left lateral prefrontal cortex and left caudate during encoding of high- vs. low-value pairs (1042 voxels at *x* = −42, *y* = 15, *z* = 30; 124 voxels at *x* = −12, *y* = −3, *z* = 9). Further, neural signatures of successful vs. unsuccessful encoding differed from those of high- vs. low-value encoding. Though we observed similar activation in left lateral PFC (1273 voxels at *x* = −48, *y* = 9, *z* = 27), here, we did not observe differential recruitment of the caudate. In addition, consistent with previous observations of subsequent memory effects (Davachi, 2006), successful encoding was associated with increased activation in the right hippocampus (21 voxels at *x* = 24, *y* = −6, *z* = −21).

### Age-related differences in neural activation during encoding

Next, we examined how neural activation during encoding vs. baseline (fixation) varied with age. Across trial types, during encoding, we observed widespread age-related increases in neural activation (Supplementary Table S31), in regions including the left lateral PFC (120 voxels at *x* = −54, *y* = 12, *z* = 33) and the right lateral PFC (24 voxels at *x* = 48, *y* = 12, *z* = 30; Figure 5C).

To address our main question of interest — how age-related differences in differential neural activation during the encoding of high- vs. low-value information may support the development of adaptive memory — we conducted two ROI analyses. Given our *a priori* hypotheses about the role of the prefrontal cortex and striatum in value-guided encoding, and their exhibiting differential activation in the high- vs. low-value encoding group-level contrast, we examined neural activation within a prefrontal cortex and striatal ROI. Despite the absence of significant differential activation in the hippocampus and parahippocampal cortex, we also used the same ROI approach to test for age differences in activation in these *a priori* regions of interest but did not observe any (See Supplement). The specific ROIs were determined by taking the peak prefrontal voxel (*x* = −51, *y* = 42, *z* = 9) and the peak striatal voxel (*x* = −18, *y* = 12, *z* = 6) from the group-level high- vs. low-value associative encoding contrast and drawing 5mm spheres around them. We then examined how the mean parameter estimate across voxels within each ROI for the high- vs. low-value encoding contrast related to both linear and quadratic age. Caudate activation did not vary significantly as a function of age (β = .16, SE = .11 *p* = .126), indicating that participants across our age range demonstrated similarly increased recruitment of the caudate while encoding high- vs. low-value associations. PFC activation, however, demonstrated a different pattern, varying as a function of both linear (β = 1.97, SE = .74, *p* = .01) and quadratic age (β = −1.73, SE = .73, *p* = .021), such that the difference in PFC engagement during encoding of high- vs. low-value associations increased to the greatest extent throughout childhood and early adolescence (Supplementary Figure S4).

The pattern of age-related differences that we observed in the PFC recruitment mirrored the age-related differences we observed in value-based memory. Given these parallel age effects across brain and behavior, we next asked whether age differences in PFC recruitment could account for our observed age differences in adaptive memory prioritization. First, we confirmed that in line with our whole-brain analysis, PFC modulation predicted memory difference scores, even when controlling for age (β = .34, SE = .10, *p* = .001). Next, we confirmed that these difference scores did in fact vary with age (β = .22, SE = .10, *p* = .041), with older participants demonstrating a larger difference in memory accuracy for high- vs. low-value associations. Critically, however, when controlling for PFC activation, age no longer related to memory difference scores (β = .15, SE = .10, *p* = .14). A formal mediation analysis revealed that PFC activation fully mediated the relation between linear age and memory difference scores (standardized indirect effect: .07, 95% confidence interval: [.01, .15], *p* = .017; standardized direct effect: .15, 95% confidence interval: [−.03, .33], *p* = .108; Figure 6). This relation was directionally specific; age did *not* mediate the relation between PFC activation and memory difference scores (standardized indirect effect: .03, 95% confidence interval: [−.007, .09], *p* = .13; standardized direct effect; .34, 95% confidence interval: [.14, .54], *p* < .001.) Further, when we included quadratic age, WASI scores, online frequency learning accuracy, online frequency learning accuracy on the final repetition of each item, and mean explicit frequency report error magnitudes as control variables in the mediation analysis, PFC activation continued to mediate the relation between linear age and memory difference scores (standardized indirect effect: .56, 95% confidence interval: [.06, 1.35], *p* = .023; standardized direct effect; 1.75, 95% confidence interval: [.12, .3.38], *p* = .034).

**Figure 6.**
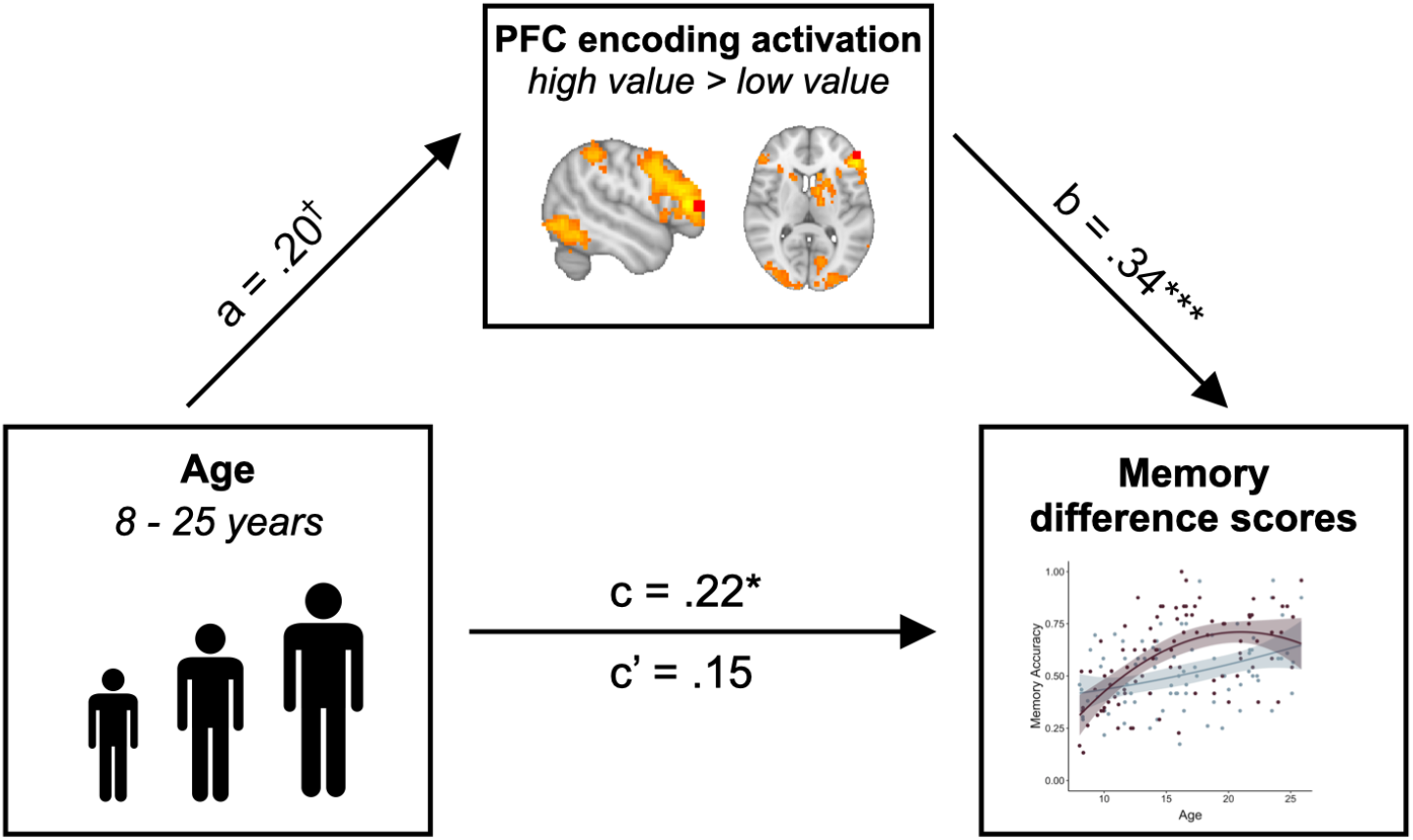
The increased engagement of left lateral PFC (ROI depicted in red) during encoding of high- vs. low-value information mediated the relation between age and memory difference scores (standardized indirect effect: .07, 95% confidence interval: [.01, .15], *p* = .017; standardized direct effect; .15, 95% confidence interval: [−.03, .33], *p* = .108). Path a shows the regression coefficient of the relation between age and PFC modulation. Path b shows the regression coefficient of the relation between PFC activation and memory difference scores, while controlling for age. Paths c and c’ show the regression coefficient of the relation between age and memory difference scores without and while controlling for PFC activation, respectively. † denotes *p* < .06, * denotes *p* < .05, ** denotes *p* < .01.

### Age- and value-based modulation of neural activation during retrieval

We next examined how the neural mechanisms of memory retrieval were influenced by both age and learned value signals. As during encoding, a whole-brain contrast comparing retrieval trials to baseline revealed age-related differences in bilateral PFC recruitment (Figure 7A; 42 voxels at *x* = 51, *y* = 3, *z* = 21; 36 voxels at *x* = −60, *y* = 6, *z* = 21) as well as regions of occipital cortex (see Supplementary Table S35) across trials during retrieval. We further tested whether participants demonstrated value-based modulation of neural activation at retrieval. During retrieval of associations involving high- vs. low-frequency items, we continued to observe increased engagement of the left lateral PFC (Figure 7B; 116 voxels at *x* = −48, *y* = 21, *z* = 24) and the bilateral caudate (128 voxels at *x* = −12, *y* = −6, *z* = 15 and 77 voxels at *x* = 15, *y* = −3, *z* = 21). This activation was not modulated by age or memory difference scores.

**Figure 7:**
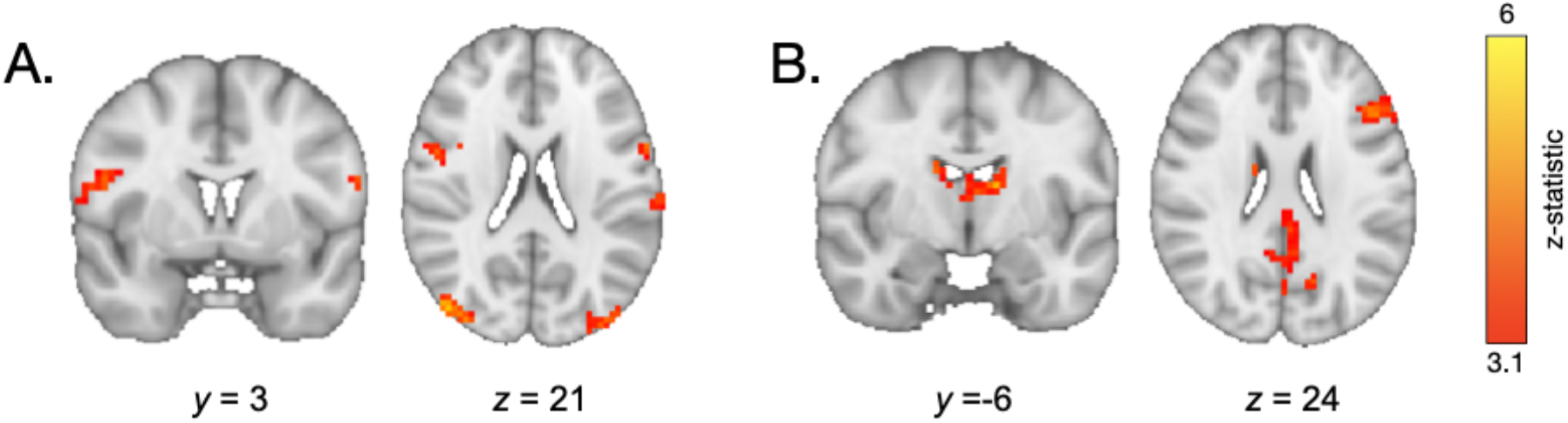
(A) During retrieval, older participants demonstrated greater recruitment of the inferior frontal cortex relative to younger participants. (B) During retrieval of associations involving high- vs. low-frequency items, participants demonstrated greater engagement of the left lateral PFC and bilateral caudate.

### Two distinct value representations influence memory

The overlap between the engagement of the neural systems we observed during encoding of high- vs. low-value information and those observed in prior studies of motivated memory that have used explicit value cues (M. S. Cohen et al., 2019, 2014) suggests that participants did indeed use learned regularities as value signals to guide memory. To what extent was memory supported by explicit representations of item frequency versus neural sensitivity to item repetitions during frequency-learning? To examine the influence of these two types of value representations across age, we ran additional mixed-effects models. First, we examined how participants’ explicit representations of item frequency related to memory. Participants demonstrated better associative memory for pairs involving items they reported were more frequent, χ^2^(1) = 31.20, *p* < .001 (Figure 8A). This effect was modulated by age (χ^2^(1) = 10.37, *p* = .001) and quadratic age (χ^2^(1) = 9.50, *p* = .002), indicating that participants’ beliefs about item frequency influenced memory to the greatest degree in adolescence and early adulthood. Further, replicating our previous behavioral findings (Nussenbaum et al., 2020), we found that the linear model including explicit frequency reports (BIC = 5438.37) fit the data *better* than the linear model including the true frequency condition (BIC = 5449.19, X^2^ = 10.83, *p* < .001), indicating that participants’ *representations* of item frequency influenced memory to a greater extent than the true item frequencies.

**Figure 8.**
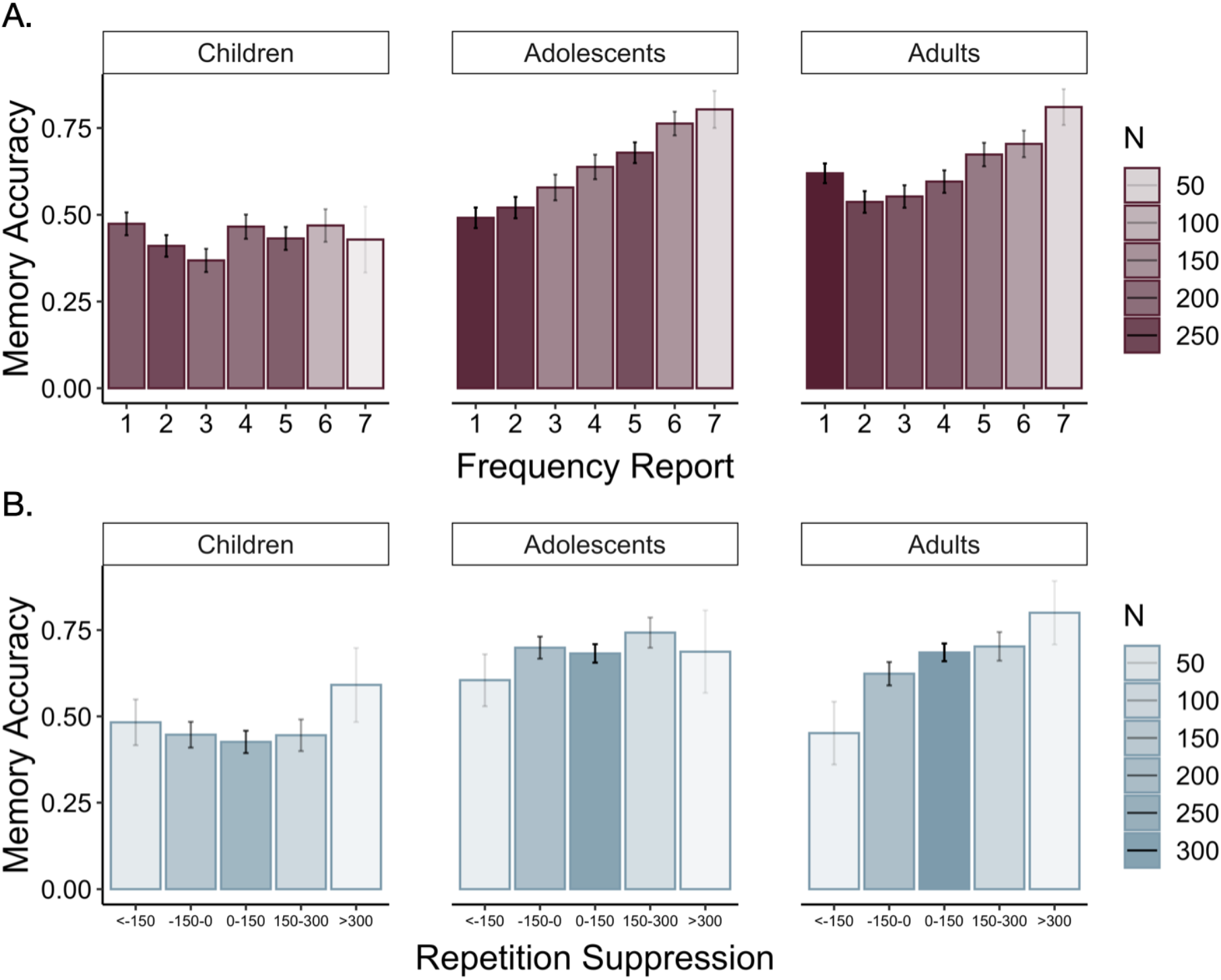
(A) Participants demonstrated increased associative memory accuracy for items that they reported as being more frequent (χ^2^(1) = 31.20, *p* < .001). This effect strengthened with increasing age (frequency report x linear age: X^2^(1) = 10.37, *p* = .001; frequency report x quadratic age: X^2^(1) = 9.50, *p* = .002). (B) Participants also demonstrated better memory for associations involving high-frequency items to which they demonstrated the greatest repetition suppression during frequency-learning (χ^2^(1) = 11.21, *p* < .001). In both panels, the shading of the bars represents the number of trials in which participants provided that frequency report.

We further examined whether our neural measure of online frequency learning related to associative memory. Specifically, we asked whether greater sensitivity to item repetitions — as indexed by greater repetition suppression within the parahippocampal cortex — promoted better encoding of associative information. Because we only had repetition suppression indices for items that appeared five times, our analysis was restricted to associations involving high-frequency items. We found that repetition suppression during frequency-learning did indeed predict subsequent associative memory, χ^2^(1) = 11.21, *p* < .001 (Figure 8B). This effect did not interact with age, χ^2^(1) = .79, *p* = .374. Further, when we included both repetition suppression indices and explicit frequency reports in our model, *both* predictors continued to explain significant variance in memory accuracy (Frequency reports: χ^2^(1) = 21.16, *p* < .001, Repetition suppression: χ^2^(1) = 10.25, *p* = .001), suggesting that learned value signals that guide memory may be derived from multiple, distinct representations of prior experience.

Given the relations we observed between memory and both repetition suppression and frequency reports, we examined whether they related to neural activation in both our caudate and PFC ROI during encoding. To do so, we computed each participant’s average repetition suppression index, and their “frequency distance” — or the average difference in their explicit reports for items in the high- and low-frequency conditions. We expected that participants with greater average repetition suppression indices and greater frequency distances represented the high- and low-frequency items as more distinct from one another and therefore would show greater differences in neural activation at encoding across frequency conditions. In line with our prior analyses, both metrics varied with age (though repetition suppression only marginally (linear age: *p* = .067; quadratic age: *p* = .042); Supplementary Tables S22 and S25), suggesting that older participants demonstrated better learning of the structure of the environment. We ran linear regressions examining the relations between each metric, age, and their interaction on neural activation in both the caudate and PFC. We observed no significant effects or interactions of average repetition suppression indices on neural activation (*p*s > .15; Supplementary Tables S23 and S24). We did, however, observe a significant effect of frequency distance on PFC activation (β = .42, SE = .12, *p* = .0012), such that participants who believed that average frequencies of the high- and low-frequency items were further apart also demonstrated greater PFC activation during encoding of pairs with high- vs. low-frequency items. Here, we did not observe a significant effect of age on PFC activation (β = −.03, SE = .13, *p* = .82), suggesting that age-related variance in PFC activation may be related to age differences in explicit frequency beliefs. Importantly, however, even when we accounted for both PFC activation and frequency distances, we continued to observe an effect of age on memory difference scores (β = .56, SE = .20, *p* = .006), which, together with our prior analyses, suggest that developmental differences in value-guided memory are not driven solely by age differences in beliefs about the structure of the environment but also depend on the use of those beliefs to guide encoding.

## Discussion

Prioritizing memory for useful information is essential throughout individuals’ lifetimes, but no prior work has investigated the development of the neural mechanisms that support value-guided memory prioritization from childhood to adulthood. The goal of the present study was to characterize how developmental differences in the neurocognitive processes that support both the *learning* and *use* of information value support improvements in adaptive memory from childhood to adulthood. In line with studies of motivated memory in adults (M. S. Cohen et al., 2014), we found that during encoding and retrieval, high-relative to low-value stimuli elicited increased activation in regions sensitive to value and motivational salience, including the caudate (Delgado et al., 2004), and those associated with strategic control processes, including the lateral PFC (Badre & Wagner, 2007; Cole & Schneider, 2007; Power & Petersen, 2013). Further, replicating previous work, we found that value-guided memory selectivity improved across childhood and adolescence (Castel et al., 2011; Hanten et al., 2007; Nussenbaum et al., 2020). Critically, here we demonstrate that increased engagement of the lateral PFC during encoding mediated the relation between age and memory selectivity. This relation was specific to the lateral PFC — though the caudate similarly demonstrated increased activation during encoding of high- vs. low-value information, value-based modulation of caudate activation did not vary as a function of age and did not relate to memory selectivity.

Two different signatures of value-learning predicted subsequent associative memory: Individuals demonstrated better memory for associations involving items that elicited stronger repetition suppression as well as for items that they reported as being more frequent. Moreover, the relation of these learning signals to memory performance varied with age. While all participants demonstrated a similar relation between repetition suppression and subsequent associative memory, the association between explicit frequency reports and memory was greater in older participants. These divergent developmental trajectories suggest that the influence of learned value on memory arises through distinct cognitive processes. One possibility is that while explicit beliefs about information value triggered the engagement of strategic control, stimulus familiarity (as indexed by repetition suppression (Gonsalves et al., 2005)) may have facilitated encoding of novel associations, even in the absence of controlled strategy use. Indeed, prior work suggests that stronger memory traces for constituent components enhances associative memory (Chalmers & Humphreys, 2003; Popov & Reder, 2020; Reder et al., 2016). However, in our previous behavioral work (Nussenbaum et al., 2020), we found that removing the relation between item frequency and reward value eliminated the memory benefit for associations involving high-frequency items, suggesting that stimulus familiarity itself did not account for the influence of item frequency on memory in our task. Still, when frequency does predict value, stimulus familiarity may serve as a proxy for information utility. This familiarity signal may exert age-invariant effects on subsequent memory, whereas explicit beliefs about item frequency may more strongly facilitate subsequent memory with increasing age.

Importantly, though we observed age-related differences in participants’ learning of the structure of their environments, the strengthening of the relation between frequency reports and associative memory with increasing age suggests that age differences in learning cannot fully account for age differences in value-guided memory. Even when accounting for individual differences in participants’ explicit knowledge of the structure of the environment, older participants demonstrated a stronger relation between their beliefs about item frequency and associative memory, suggesting that they used their beliefs to guide memory to a greater degree than younger participants. In addition, we continued to observe a robust interaction between age and frequency condition on associative memory, even when controlling for age-related change in the accuracy of both online frequency learning and explicit frequency reports. Thus, though we observed age differences in the learning of environmental regularities *and* in their influence on subsequent associative memory encoding, our developmental memory effects cannot be fully explained by differences in initial learning.

Our neural results further suggest that developmental differences in memory were driven by both knowledge of the structure of the environment and *use* of that knowledge to guide encoding. Specifically, we observed age-related increases in both overall PFC engagement as well as its value-based modulation, which may reflect developmental changes in the engagement of strategic control. Our finding that lateral prefrontal cortex activation during encoding of high- vs. low-value information may underpin memory selectivity is also in line with prior studies of motivated memory in *older* adults (M. S. Cohen et al., 2016). Older adults have been shown to demonstrate decreased neural activation in response to value cues (M. S. Cohen et al., 2016; Geddes et al., 2018) but preserved memory selectivity (Castel, 2007; Castel et al., 2002; M. S. Cohen et al., 2016) supported by the strategic recruitment of the left lateral PFC during encoding of high- vs. low-value information (M. S. Cohen et al., 2016). The PFC may support enhanced attention (Uncapher et al., 2011; Uncapher & Wagner, 2009) and semantic elaboration (Kirchhoff & Buckner, 2006) during encoding, and more focused (Wais et al., 2012) and organized search and selection (Badre & Wagner, 2007; Yu et al., 2018) during retrieval. From childhood to young adulthood, individuals demonstrate improvements in the implementation of strategic memory processes (Bjorkland et al., 2009; Yu et al., 2018), which are paralleled by increases in PFC recruitment during both encoding and retrieval (Ghetti et al., 2010; Ghetti & Fandakova, 2020; Ofen et al., 2007; Shing et al., 2016; Tang et al., 2018). In line with this prior work, we similarly observed age-related improvements in overall memory performance and in prefrontal recruitment during encoding and retrieval of novel associations.

The development of adaptive memory requires not only the implementation of encoding and retrieval strategies, but also the flexibility to up- or down-regulate the engagement of control in response to momentary fluctuations in information value (Castel et al., 2007, 2013; Hennessee et al., 2017). Importantly, value-based modulation of lateral PFC engagement during encoding mediated the relation between age and memory selectivity, suggesting that developmental change in both the representation of learned value and value-guided cognitive control may underpin the emergence of adaptive memory prioritization. Prior work examining other neurocognitive processes, including response inhibition (Insel et al., 2017) and selective attention (Störmer et al., 2014), has similarly found that increases in the flexible upregulation of control in response to value cues enhance goal-directed behavior across development (Davidow et al., 2018), and may depend on the engagement of both striatal and prefrontal circuitry (Hallquist et al., 2018; Insel et al., 2017). Here, we extend these past findings to the domain of memory, demonstrating that value signals derived from the structure of the environment increasingly elicit prefrontal cortex engagement and strengthen goal-directed encoding across childhood and into adolescence.

Further, we also demonstrate that in the absence of *explicit* value cues, the engagement of prefrontal control processes may reflect *beliefs* about information value that are learned through experience. Here, we found that differential PFC activation during encoding of high- vs. low-value information reflected individual and age-related differences in beliefs about the structure of the environment; participants who represented the average frequencies of the low- and high-frequency items as further apart *also* demonstrated greater value-based modulation of lateral PFC activation. It is important to note, however, that we collected explicit frequency reports *after* associative encoding and retrieval. Thus, the relation between PFC activation and explicit frequency reports may be bidirectional — while participants may have increased the recruitment of cognitive control processes to better encode information they believed was more valuable, the engagement of more elaborative or deeper encoding strategies that led to stronger memory traces may have also increased participants’ subjective sense of an item’s frequency (Jonides & Naveh-Benjamin, 1987).

During retrieval, we continued to observe increased activation of the caudate and dlPFC for high- vs. low-value pairs. However, this activation did not significantly vary as a function of memory difference scores or age, suggesting that the developmental differences in value-guided memory that we observed were likely driven by age-related change in encoding processes.

We found that memory prioritization varied with *quadratic* age, and our follow-up tests probing the quadratic age effect did not reveal evidence for significant age-related change in memory prioritization between late adolescence and early adulthood. However, in our prior behavioral work using a very similar paradigm (Nussenbaum et al., 2020), we found that memory prioritization varied with *linear* age only. In line with theoretical proposals (Davidow et al., 2018), subtle differences in the control demands between the two tasks (e.g., reducing the number of ‘foils’ presented on each trial of the memory test here relative to our prior study), may have shifted the age range across which we observed differences in behavior, with the more demanding variant of our task showing more linear age-related improvements into early adulthood. In addition, the specific control demands of our task may have also influenced the age at which value-guided memory emerged. Future studies should test whether younger children can modulate encoding based on the value of information if the mnemonic demands of the task are simpler.

One important caveat is that our study was cross-sectional — it will be important to replicate our findings in a longitudinal sample to measure more directly how developmental *changes* in cognitive control within an individual contribute to changes in their ability to selectively encode useful information. Our mediation results, in particular, must be interpreted with caution, as simulations have demonstrated that in cross-sectional samples, variables can emerge as significant mediators of age-related change due largely to statistical artifact (Hofer et al., 2006; Lindenberger et al., 2011). Indeed, our finding that PFC activation mediates the relation between age and value-guided memory does not necessarily imply that within an individual, PFC development leads to improvements in memory selectivity. Longitudinal work in which individuals’ neural activity and memory performance is sampled densely within developmental windows of interest is needed to elucidate the complex relations between age, brain development, and behavior (Hofer et al., 2006; Lindenberger et al., 2011).

We did not find evidence to support two of our predictions. First, though we initially hypothesized that both the ventral and dorsal striatum may be involved in encoding of high-value information, the activation we observed was largely within the dorsal striatum, a region that may reflect the value of goal-directed actions (Liljeholm & O’Doherty, 2012). In our task, rather than each stimulus acquiring intrinsic value during frequency-learning, participants may have represented the value of the ‘action’ of remembering each pair during encoding. Second, we did not observe differences in hippocampal engagement during encoding or retrieval of high- vs. low-value associates. This is somewhat surprising, as prior work has suggested that value cues bolster memory through their influence on medial temporal lobe activity (Adcock et al., 2006). However, because our task required the use of *learned* value signals, memory retrieval processes may have been required on every encoding trial to recall the frequency of each item. Thus, all items presented at encoding may have triggered the engagement of hippocampal-dependent retrieval processes (Squire, 1992) that underpin many forms of memory-guided behavior including attention (Stokes et al., 2012; Summerfield et al., 2006) and decision-making (Murty et al., 2016; Shadlen & Shohamy, 2016; Wang et al., 2020). Further, in line with prior work (Davachi, 2006), we did observe increased activation in the hippocampus during encoding of associations that were subsequently remembered vs. those that were subsequently forgotten. Medial temporal activation, including hippocampal activation, may thus more strongly reflect successful memory formation — whether or not it was facilitated by value — whereas the caudate and lateral prefrontal cortex may be more sensitive to fluctuations in the engagement of value-guided cognitive control.

There are multiple routes through which value signals influence memory (M. S. Cohen et al., 2019), and in many contexts, reward-motivated memory may not require strategic control. Value anticipation and reward delivery lead to dopaminergic release in the VTA, which projects not only to corticostriatal circuits that implement goal-directed strategy selection (Liljeholm & O’Doherty, 2012), but also directly to the hippocampus and medial temporal lobes (Adcock et al., 2006; Lisman & Grace, 2005; Murty et al., 2017; Shohamy & Adcock, 2010; Stanek et al., 2019). Given the earlier development of subcortically restricted circuitry relative to the more protracted development of cortical-subcortical pathways (Somerville & Casey, 2010), it may be the case that the *direct* influence of reward on memory develops earlier than the controlled pathway we studied here. Rather than eliciting strategic control through incentivizing successful memory, this pathway can be engaged through direct delivery of rewards or reinforcement signals at the time of encoding (Ergo et al., 2020; Jang et al., 2019; Rosenbaum et al., 2020; Rouhani et al., 2018). In one such study, adolescents demonstrated greater reward-based modulation of hippocampal-striatal connectivity than adults, and the strength of this connectivity predicted reward-related memory (Davidow et al., 2016). However, other studies have not found evidence for developmental change in the influence of valenced outcomes on memory (A. O. Cohen et al., 2019; Katzman & Hartley, 2020). The influence of different motivational and reward signals on memory across development may not be straightforward — individual and developmental differences in neurocognitive processes including sensitivity to valenced feedback (Ngo et al., 2019; Rosenbaum et al., 2020), curiosity (Fandakova & Gruber, 2021), and emotional processing (Adelman & Estes, 2013; Eich & Castel, 2016) may interact, leading to complex relations between age, motivation, and memory performance. Further work is needed to characterize both the influence of different types of reward signals on memory across development, as well as the development of the neural pathways that underlie age-related change in behavior.

The present study contributes to our understanding of the neurocognitive mechanisms that support memory across development. Specifically, we addressed the question of how motivated memory may operate in the absence of explicit value cues by examining the development of the neurocognitive mechanisms that support the *learning* and *use* of information value to guide encoding and retrieval. The present findings suggest that while development is marked by improvements in the ability to learn about the statistical structure of the environment, the emergence of adaptive memory also depends centrally on age-related differences in prefrontal control. Our findings demonstrate that prefrontal cortex development has implications not just for general memory processes but for the selective prioritization of useful information — a key component of adaptive memory throughout the lifespan.

## Methods

### Participants

Ninety participants between the ages of 8.0 and 25.9 years took part in this experiment. Thirty participants (n = 16 females) were children between the ages of 8.0 and 12.7 years, 30 participants were adolescents between the ages of 13.0 and 17.7 years (n = 16 females), and 30 participants were adults between the ages of 18.32 and 25.9 years (n = 15 females). Ten additional participants were tested but excluded from all analyses due to excessive motion during the fMRI scan (n = 8; see exclusion criteria below) or technical errors during data acquisition (n = 2). We based our sample size on other functional neuroimaging studies of the development of goal-directed behavior and memory across childhood and adolescence (Insel et al., 2017; Tang et al., 2018) as well as on our prior behavioral study that showed age-related change in the use of learned value to guide memory (Nussenbaum et al., 2020). According to self- or parental-report, participants were right-handed, had normal or corrected-to-normal vision, and no history of diagnosed psychiatric or learning disorders. Participants were recruited via flyers around New York University, and from science fairs and events throughout New York City. Based on self- or parent-report, 35.6% of participants were White, 26.7% were two or more races, 24.4% were Asian, 11.1% were Black and 2.2% were Native American. Additionally, 17.8% of the sample identified as Hispanic.

Research procedures were approved by New York University’s Institutional Review Board. Adult participants provided written consent prior to participating in the study. Children and adolescents provided written assent, and their parents or guardians provided written consent on their behalf, prior to their participation. All participants were compensated $60 for the experimental session, which involved a 1-hour MRI scan. Participants were told that they would receive an additional bonus payment based on their performance in the experiment; in reality, all participants received an additional $5 bonus payment.

Prior to participating in the scanning session, child and adolescent participants who had never participated in a MRI study in our lab completed a mock scanning session to acclimate to the scanning environment. Mock scan sessions took place during a separate lab visit, at least one day in advance of scheduled scans. In the mock scanner, participants practiced staying as still as possible. We attached a Wii-mote to their heads, and set it to “rumble” whenever it sensed that the participant had moved. Participants completed a series of three challenges of increasing duration (10, 30, and 90 seconds) and decreasing angular tolerance (10, 5, and 2 degrees) in which they tried to prevent the Wii-mote from rumbling by lying very still (Casey et al., 2018).

### Experimental Tasks

Participants completed two blocks of three tasks (Figure 1), a variant of which we used in a previous behavioral study (Nussenbaum et al., 2020). Across tasks, participants made responses with two MRI-compatible button boxes, one for each hand. In between tasks, an experimenter reminded participants of the instructions for the next part, and participants viewed a diagram indicating which fingers and buttons they should use to make their responses. The tasks were presented using Psychtoolbox Version 3 (Brainard, 1997; Kleiner et al., 2007; Pelli, 1997) for Matlab 2017a (Mathworks Inc, 2017) and displayed on a screen behind the scanner, visible to participants via a mirror attached to the MRI head coil. FMRI BOLD activity was measured over eight functional runs, which ranged in duration from approximately 4 to 7.5 minutes.

The structure of each block of tasks was identical, but their narratives and stimuli differed. In one set of tasks, participants were told that they had a collection of postcards they needed to mail. Each type of postcard in their collection required a different type of stamp.

In the *frequency-learning* task, participants were told they had to sort through their postcards to learn how many of each type they had. They were told that they had more of some types of postcards relative to other types (e.g., they might have 5 postcards with the same, specific blue pattern but only 1 postcard with a specific red pattern (Figure 1)). Participants were instructed to try to keep track of how many of each kind of postcard they had, because it would be useful to them later on. Throughout the task, participants viewed 24 images of postcards. Twelve of these images were presented once and 12 of the images were presented five times, such that participants completed 72 trials total. On each trial, a postcard appeared in the center of the screen for 2 seconds. Across all tasks, stimulus presentation was followed by an inter-trial interval (ITI) of 2 - 6 seconds, which consisted of a black screen with a small, white fixation cross. Participants were instructed to press the button under their right index finger when they saw a new postcard they had not seen before and to press the button under their right middle finger when they saw a repeated postcard that they had already seen within the task. Participants were instructed to respond as quickly and as accurately as possible. The specific postcard assigned to each frequency condition (1 or 5) was counterbalanced across participants. The order of image presentation was randomized for each participant.

In the second task, the *associative encoding* task, participants were told that they would learn the correct stamp to put on each type of postcard. Participants were instructed that in the subsequent task, they would have to stamp all of their postcards, earning one point for each postcard stamped correctly. Critically, in the associative encoding task, regardless of the number of each type of postcard that they had (i.e., 1 or 5), participants saw each type of postcard with its corresponding stamp only *once*. Participants were instructed that they would earn more points if they focused on remembering the stamps that went on the types of postcards that they had the most of. Thus participants had equal exposure to the to-be-encoded associations across frequency conditions. On each trial, participants viewed one of the types of postcards from the frequency task next to an image of a unique stamp (5 s). The stamp-postcard pairs, order of the trials, and side of the screen on which the stamp and postcard appeared were randomized for each participant.

Next, participants completed *retrieval*. In the first part of the retrieval task, participants viewed all 24 unique postcards, one at time. When each postcard appeared, participants also saw four stamps: the correct stamp, a foil stamp that had been presented with a high-frequency postcard in the previous paired-associates task, a foil stamp that had been presented with a low-frequency postcard, and a novel stamp. Participants used the four fingers on their right hands to select one of these four stamps. Participants had six seconds to make their selection. Regardless of when they made their selection, the card and all four stamps remained on the screen for 6 seconds. After participants pressed a stamp, a faint, grey outline appeared around it. No feedback was given until the end of the set of tasks. The order of the postcard and the location of each stamp was randomized for each participant.

After stamping all 24 unique postcards once, participants’ memory for the postcards’ original frequencies was then probed. Participants again saw all 24 unique postcards, one at a time this time with the numbers 1 – 7 underneath them, and they were asked to provide *frequency reports*. Participants used three fingers on their left hand and all four fingers on their right hand to select the number that they believed matched the number of times they saw the card in the first task. As in the previous task, participants had six seconds to make their selection. Regardless of when they made their selection, the card and all seven numbers remained on the screen for 6 seconds. After participants selected a number, a faint, grey outline appeared around it. The order of the postcards was randomized for each participant.

Finally, participants stamped all remaining postcards, such that they completed 48 additional memory test trials (i.e. they stamped each of the postcards in the 5-frequency condition four more times.) These trials were not included in any analyses, but their inclusion ensured that correctly encoding the stamps that belonged on the high-frequency postcards would be more valuable for participants despite each retrieval trial being worth 1 point. Here, participants had 4 seconds to make each response. We did not record neural data for this run, so each trial was followed by a 500-ms black screen with a white fixation cross. At the end of the memory test, participants saw a screen that displayed how many postcards they stamped correctly.

After completing the three tasks, participants were told that they were going to play a second set of similar games. The second set of tasks was identical to the first, except that the stimuli were changed from postcards and stamps to landscape pictures and picture frames. The order of the stimulus sets was counterbalanced across participants.

Prior to entering the scanner, all participants completed a short task tutorial on a laptop to learn the overall task structure and the instructions for each part. The task tutorial comprised a full set of identical tasks but with only two stimuli within each frequency condition. Participants who first did the tasks with postcards completed a tutorial with four novel postcards and novel stamps; participants who first did the tasks with pictures completed a tutorial with four novel pictures and novel frames.

Child and adolescent participants were administered the Vocabulary and Matrix Reasoning subtests of the Wechsler Abbreviated Scale of Intelligence (WASI) (Wechsler, 2011) during the mock scanning session. Adults were administered the same two subtests immediately following their scan. We followed the standard procedure to compute age-normed IQ scores for each participant based on their performance on these two sub-tests.

### Analysis of behavioral data

All behavioral data processing and statistical analyses were conducted in R version 3.5.1 (R Core Team, 2018). Data were combined across blocks (but we include an analysis of block effects on memory performance in the supplement). Trials in which participants failed to make a response were excluded from analyses. Mixed effects models were run using the ‘afex’ package version 0.21-2 (Singmann et al., 2020). Numeric variables were *z*-scored across the entire data set prior to their inclusion in each model. To determine the random effects structures of our mixed effects models, we began with the maximal model to minimize Type I errors (Barr et al., 2013). We included random participant intercepts and slopes across all fixed effects (except age and WASI scores) and their interactions. We also included random stimulus intercepts and slopes across all fixed effects and their interactions. Because stimuli were randomly paired during associative encoding and only repeated, on average, around four times across participants, our stimulus random effects accounted for individual items (e.g., postcard 1) rather than pairs of items (e.g., postcard 1 and stamp 5). We set the number of model iterations to one million and use the “bobyqa” optimizer. When the maximal model gave convergence errors or failed to converge within a reasonable timeframe (~24 hours), we removed correlations between random slopes and random intercepts, followed by random slopes for interaction effects, followed by random slopes across stimuli. For full details about the fixed- and random-effects structure of all models, see “Full Model Specification and Results” in the supplement. To test the significance of the fixed effects in our models, we used likelihood ratio tests for logistic models and F tests with Satterthwaite approximations for degrees of freedom for linear models. Mediation analyses were conducted with the “mediation” R package (Tingley et al., 2014) and significance of the mediation effects was assessed via 10,000 bootstrapped samples.

For our memory analyses, trials were scored as ‘correct’ if the participant selected the correct association from the set of four possible options presented during the memory test, ‘incorrect’ if the participant selected an incorrect association, and ‘missed’ if the participant failed to respond within the 6-second response window. Missed trials were excluded from all analyses. Because participants had to select the correct association from four possible options, chance-level performance was 25%. Two child participants performed at or below chance-level on the memory test. They were included in all analyses reported in the manuscript; however, we report full details of the results of our memory analyses when we exclude these two participants in the supplement (Supplementary Table S15). Importantly, our main findings remain unchanged.

### Image acquisition, preprocessing, and quality assessment

Participants were scanned at New York University’s Center for Brain Imaging using a Siemens Prisma 3T MRI scanner with a 64-channel head coil. Anatomical data were acquired with high-resolution, T1- weighted anatomical scans using a magnetization-prepared rapidly acquired gradient echo (MPRAGE) sequence (TR = 2.3s, TE = 2.3ms, TI = .9s; 8° flip angle; .9-mm isotropic voxels, field of view = 192 × 256 × 256 voxels; acceleration: GRAPPA 2 in the phase-encoding direction, with 24 reference lines) and T2- weighted anatomical scans using a 3D turbo spin echo (TSE) sequence (T2: TR = 3.2s, TE = 564ms, Echo Train Length = 314; 120° flip angle, .9-mm isotropic voxels, field of view = 240 × 256 × 256 voxels; acceleration: GRAPPA 2×2 with 32 reference lines in both the phase- and slice-encoding directions). Functional data were acquired with a T2*-weighted, multi-echo EPI sequence with the following parameters: TR=2s, TEs=12.2, 29.48, 46.76, 64.04ms; MB factor = 2; acceleration: GRAPPA 2, with 24 reference lines; effective echo spacing: .245 ms; 44 axial slices; 75° flip angle, 3-mm isotropic voxels, from the University of Minnesota’s Center for Magnetic Resonance Research (Feinberg et al., 2010; Moeller et al., 2010; Xu et al., 2013).

All anatomical and functional MRI data were preprocessed using fMRIPrep v.1.5.1rc2 (Esteban et al., 2019), a robust preprocessing pipeline that adjusts to create the optimal workflow for the input dataset, and then visually inspected. FMRIPrep uses tedana (for implementation details, see (Kundu et al., 2013, 2012) to combine each four-echo time series based on the signal decay rate of each voxel, taking a weighted average of the four echoes that optimally balances signal strength and BOLD sensitivity. This approach enables the acquisition of BOLD data with a higher signal-to-noise ratio, giving us greater sensitivity to detect neural effects of interest (Kundu et al., 2013). This combined time series is then used in subsequent preprocessing steps (e.g., susceptibility distortion correction, confound estimation, registration). Runs in which more than 15% of TRs were censored for motion (relative motion > .9 mm framewise displacement) were excluded from neuroimaging analyses (see Supplementary Table S1 for the number of participants included in each analysis). Participants who did not have at least one usable run of each task (frequency-learning, associative encoding, retrieval), were excluded from all behavioral and neuroimaging analyses (n = 8), leaving N = 90 participants in our analyzed sample.

### Analysis of fMRI data

Statistical analyses were completed in FSL v. 6.0.2. (Jenkinson et al., 2012; Smith et al., 2004). Preprocessed BOLD data, registered to fMRIPrep’s MNI152 template space and smoothed with a 5mm Gaussian kernel, were combined across runs via fixed-effects analyses and then submitted to mixed-effects GLM analyses, implemented in FEAT 6.0.0 (M. W. Woolrich et al., 2001; Mark W. Woolrich et al., 2004), to estimate relevant task effects. For all GLM analyses, nuisance regressors included six motion parameters and their derivatives, framewise displacement values, censored frames, the first six anatomical noise components (aCompCor) from fMRIPrep, and cosine regressors from fMRIprep to perform high-pass filtering of the data. All task-based temporal onset regressors were convolved with a double gamma hemodynamic response function and included temporal derivatives. Analyses were thresholded using a wholebrain correction of *z* > 3.1 and a cluster-defining threshold of *p* < .05 using FLAME 1.

#### Frequency-learning GLM

Our frequency-learning model included six task-based temporal onset regressors. Trials were divided based on appearance count and frequency condition to create the following regressors: 1) Low-frequency items the first (and only) time they appeared, 2) High-frequency items the first time they appeared, 3) High-frequency items the second time they appeared, 4) High-frequency items the third time they appeared, 5) High-frequency items the fourth time they appeared, 6) High-frequency items the fifth time they appeared.

#### Repetition suppression analyses

For each stimulus in the high-frequency condition, we examined repetition suppression by measuring activation within a parahippocampal ROI during the presentation of each item during frequency-learning. We defined our ROI by taking the peak voxel (*x* = 30, *y* = −39, *z* = −15) from the group-level first > last item appearance contrast for high-frequency items during frequency-learning and drawing a 5 mm sphere around it. This voxel was located in the right parahippocampal cortex, though we observed widespread and largely symmetric activation in bilateral parahippocampal cortex. To encompass both left and right parahippocampal cortex within our ROI, we mirrored the peak voxel sphere. For each participant, we modeled the neural response to each appearance of each item using the Least Squares-Separate approach (Mumford et al., 2014). Each first-level model included a regressor for the trial of interest, as well as separate regressors for the onsets of all other items, grouped by repetition number (e.g., a regressor for item onsets on their first appearance, a regressor for item onsets on their second appearance, etc.). Values that fell outside five standard deviations from the mean level of neural activation across all subjects and repetitions were excluded from subsequent analyses (18 out of 10,320 values; .01% of observations). In addition to examining neural activation as a function of stimulus repetition, we also computed an index of repetition suppression for each high-frequency item by computing the difference in mean beta values within our ROI on its first and last appearance.

#### Associative encoding and retrieval GLMs

Our associative encoding and retrieval models included six task-based temporal onset regressors. Trials were divided based on frequency condition (high- vs. low-) and subsequent memory (remembered, forgotten, missed). Missed trials were included as nuisance regressors and not included in any contrasts.

#### Associative encoding regions of interest (ROIs)

Given our *a priori* hypotheses about the role of the prefrontal cortex and striatum in value-guided encoding, we examined neural activation within a prefrontal cortex and striatal ROI. The specific ROIs were determined by taking the peak prefrontal voxel (*x* = −51, *y* = 42, *z* = 9) and the peak striatal voxel (*x* = −18, *y* = 12, *z* = 6) from the group-level high > low value associative encoding contrast and drawing 5mm spheres around them.

## Acknowledgments

We thank Daphne Valencia, Jamie Greer, Nora Keathley, and Michael Liu for assistance with data collection, Ali Cohen and Gail Rosenbaum for valuable discussions and help with analyses, and the staff of NYU’s Center for Brain Imaging, especially Pablo Velasco, for technical support and guidance. This project was supported by a National Science Foundation CAREER Grant (1654393) to C.A.H., a Jacobs Foundation Early Career Fellowship to C.A.H. and a National Defense Science and Engineering Graduate Fellowship to K.N.

## Data and Code Availability

Task code, behavioral data, and analysis code are available on the Open Science Framework: https://osf.io/2fkbi/. Unthresholded *z*-statistic maps from the neuroimaging analyses are available on Neurovault: https://neurovault.org/collections/BUMNZQXA/. Upon publication, BIDS-formatted neuroimaging data will be made public on Open Neuro: https://openneuro.org/datasets/ds003499

## References

Adcock, R. A., Thangavel, A., Whitfield-Gabrieli, S., Knutson, B., & Gabrieli, J. D. E. (2006). Reward-motivated learning: mesolimbic activation precedes memory formation. Neuron, 50(3), 507–517. https://doi.org/10.1016/j.neuron.2006.03.036

Adelman, J. S., & Estes, Z. (2013). Emotion and memory: a recognition advantage for positive and negative words independent of arousal. Cognition, 129(3), 530–535. https://doi.org/10.1016/j.cognition.2013.08.014

Amso, D., & Davidow, J. (2012). The development of implicit learning from infancy to adulthood: item frequencies, relations, and cognitive flexibility. Developmental Psychobiology, 54(6), 664–673. https://doi.org/10.1002/dev.20587

Anderson, J. R., & Milson, R. (1989). Human memory: An adaptive perspective. In Psychological Review (Vol. 96, Issue 4, pp. 703–719). https://doi.org/10.1037//0033-295x.96.4.703

Anderson, J. R., & Schooler, L. J. (1991). Reflections of the Environment in Memory. Psychological Science, 2(6), 396–408. https://doi.org/10.1111/j.1467-9280.1991.tb00174.x

Auksztulewicz, R., & Friston, K. (2016). Repetition suppression and its contextual determinants in predictive coding. Cortex; a Journal Devoted to the Study of the Nervous System and Behavior, 80, 125–140. https://doi.org/10.1016/j.cortex.2015.11.024

Badre, D., & Wagner, A. D. (2007). Left ventrolateral prefrontal cortex and the cognitive control of memory. Neuropsychologia, 45(13), 2883–2901. https://doi.org/10.1016/j.neuropsychologia.2007.06.015

Barr, D. J., Levy, R., Scheepers, C., & Tily, H. J. (2013). Random effects structure for confirmatory hypothesis testing: Keep it maximal. Journal of Memory and Language, 68(3). https://doi.org/10.1016/j.jml.2012.11.001

Biderman, N., Bakkour, A., & Shohamy, D. (2020). What Are Memories For? The Hippocampus Bridges Past Experience with Future Decisions. Trends in Cognitive Sciences, 24(7), 542–556. https://doi.org/10.1016/j.tics.2020.04.004

Bjorkland, D. F., Dukes, C., & Brown, D. R. (2009). The development of memory strategies. The Development of Memory in Infancy and Childhood. Psychology Press, NY, 145–176.

Botvinick, M., & Braver, T. (2015). Motivation and cognitive control: from behavior to neural mechanism. Annual Review of Psychology, 66, 83–113. https://doi.org/10.1146/annurev-psych-010814-015044

Braams, B. R., van Duijvenvoorde, A. C. K., Peper, J. S., & Crone, E. A. (2015). Longitudinal changes in adolescent risk-taking: a comprehensive study of neural responses to rewards, pubertal development, and risk-taking behavior. The Journal of Neuroscience: The Official Journal of the Society for Neuroscience, 35(18), 7226–7238. https://doi.org/10.1523/JNEUROSCI.4764-14.2015

Brainard, D. H. (1997). The Psychophysics Toolbox. Spatial Vision, 10(4), 433–436. https://www.ncbi.nlm.nih.gov/pubmed/9176952

Casey, B. J., Cannonier, T., Conley, M. I., Cohen, A. O., Barch, D. M., Heitzeg, M. M., Soules, M. E., Teslovich, T., Dellarco, D. V., Garavan, H., Orr, C. A., Wager, T. D., Banich, M. T., Speer, N. K., Sutherland, M. T., Riedel, M. C., Dick, A. S., Bjork, J. M., Thomas, K. M., … ABCD Imaging Acquisition Workgroup. (2018). The Adolescent Brain Cognitive Development (ABCD) study: Imaging acquisition across 21 sites. Developmental Cognitive Neuroscience, 32, 43–54. https://doi.org/10.1016/j.dcn.2018.03.001

Castel, A. D. (2007). The Adaptive and Strategic Use of Memory By Older Adults: Evaluative Processing and Value-Directed Remembering. In Psychology of Learning and Motivation(pp. 225–270). https://doi.org/10.1016/s0079-7421(07)48006-9

Castel, A. D., Benjamin, A. S., Craik, F. I. M., & Watkins, M. J. (2002). The effects of aging on selectivity and control in short-term recall. Memory & Cognition, 30(7), 1078–1085. https://doi.org/10.3758/bf03194325

Castel, A. D., Farb, N. A. S., & Craik, F. I. M. (2007). Memory for general and specific value information in younger and older adults: measuring the limits of strategic control. Memory & Cognition, 35(4), 689–700. https://www.ncbi.nlm.nih.gov/pubmed/17848027

Castel, A. D., Humphreys, K. L., Lee, S. S., Galván, A., Balota, D. A., & McCabe, D. P. (2011). The development of memory efficiency and value-directed remembering across the life span: a cross-sectional study of memory and selectivity. Developmental Psychology, 47(6), 1553–1564. https://doi.org/10.1037/a0025623

Castel, A. D., Murayama, K., Friedman, M. C., McGillivray, S., & Link, I. (2013). Selecting valuable information to remember: age-related differences and similarities in self-regulated learning. Psychology and Aging, 28(1), 232–242. https://doi.org/10.1037/a0030678

Chalmers, K. A., & Humphreys, M. S. (2003). Experimental manipulation of prior experience: Effects on item and associative recognition. Memory (Hove, England), 11(3), 233–246. https://doi.org/10.1080/09658210244000009

Cohen, A. O., Matese, N. G., Filimontseva, A., Shen, X., Shi, T. C., Livne, E., & Hartley, C. A. (2019). Aversive learning strengthens episodic memory in both adolescents and adults. In Learning & Memory (Vol. 26, Issue 7, pp. 272–279). https://doi.org/10.1101/lm.048413.118

Cohen, M. S., Cheng, L. Y., Paller, K. A., & Reber, P. J. (2019). Separate Memory-Enhancing Effects of Reward and Strategic Encoding. Journal of Cognitive Neuroscience, 31(11), 1658–1673. https://doi.org/10.1162/jocn_a_01438

Cohen, M. S., Rissman, J., Suthana, N. A., Castel, A. D., & Knowlton, B. J. (2014). Value-based modulation of memory encoding involves strategic engagement of fronto-temporal semantic processing regions. Cognitive, Affective & Behavioral Neuroscience, 14(2), 578–592. https://doi.org/10.3758/s13415-014-0275-x

Cohen, M. S., Rissman, J., Suthana, N. A., Castel, A. D., & Knowlton, B. J. (2016). Effects of aging on value-directed modulation of semantic network activity during verbal learning. NeuroImage, 125, 1046–1062. https://doi.org/10.1016/j.neuroimage.2015.07.079

Cole, M. W., & Schneider, W. (2007). The cognitive control network: Integrated cortical regions with dissociable functions. NeuroImage, 37(1), 343–360. https://doi.org/10.1016/j.neuroimage.2007.03.071

Davachi, L. (2006). Item, context and relational episodic encoding in humans. In Current Opinion in Neurobiology (Vol. 16, Issue 6, pp. 693–700). https://doi.org/10.1016/j.conb.2006.10.012

Davidow, J. Y., Foerde, K., Galván, A., & Shohamy, D. (2016). An Upside to Reward Sensitivity: The Hippocampus Supports Enhanced Reinforcement Learning in Adolescence. Neuron, 92(1), 93–99. https://doi.org/10.1016/j.neuron.2016.08.031

Davidow, J. Y., Insel, C., & Somerville, L. H. (2018). Adolescent Development of Value-Guided Goal Pursuit. Trends in Cognitive Sciences, 22(8), 725–736. https://doi.org/10.1016/j.tics.2018.05.003

Delgado, M. R., Stenger, V. A., & Fiez, J. A. (2004). Motivation-dependent responses in the human caudate nucleus. Cerebral Cortex, 14(9), 1022–1030. https://doi.org/10.1093/cercor/bhh062

Eich, T. S., & Castel, A. D. (2016). The cognitive control of emotional versus value-based information in younger and older adults. Psychology and Aging, 31(5), 503–512. https://doi.org/10.1037/pag0000106

Ergo, K., De Loof, E., & Verguts, T. (2020). Reward Prediction Error and Declarative Memory. Trends in Cognitive Sciences, 24(5), 388–397. https://doi.org/10.1016/j.tics.2020.02.009

Esteban, O., Markiewicz, C. J., Blair, R. W., Moodie, C. A., Isik, A. I., Erramuzpe, A., Kent, J. D., Goncalves, M., DuPre, E., Snyder, M., Oya, H., Ghosh, S. S., Wright, J., Durnez, J., Poldrack, R. A., & Gorgolewski, K. J. (2019). fMRIPrep: a robust preprocessing pipeline for functional MRI. Nature Methods, 16(1), 111–116. https://doi.org/10.1038/s41592-018-0235-4

Fandakova, Y., & Gruber, M. J. (2021). States of curiosity and interest enhance memory differently in adolescents and in children. Developmental Science, 24(1), e13005. https://doi.org/10.1111/desc.13005

Feinberg, D. A., Moeller, S., Smith, S. M., Auerbach, E., Ramanna, S., Glasser, M. F., Miller, K. L., Ugurbil, K., & Yacoub, E. (2010). Multiplexed Echo Planar Imaging for Sub-Second Whole Brain FMRI and Fast Diffusion Imaging. In PLoS ONE (Vol. 5, Issue 12, p. e15710). https://doi.org/10.1371/journal.pone.0015710

Finn, A. S., Kalra, P. B., Goetz, C., Leonard, J. A., Sheridan, M. A., & Gabrieli, J. D. E. (2016). Developmental dissociation between the maturation of procedural memory and declarative memory. Journal of Experimental Child Psychology, 142, 212–220. https://doi.org/10.1016/j.jecp.2015.09.027

Geddes, M. R., Mattfeld, A. T., Angeles, C. de L., Keshavan, A., & Gabrieli, J. D. E. (2018). Human aging reduces the neurobehavioral influence of motivation on episodic memory. NeuroImage, 171, 296–310. https://doi.org/10.1016/j.neuroimage.2017.12.053

Ghetti, S., DeMaster, D. M., Yonelinas, A. P., & Bunge, S. A. (2010). Developmental differences in medial temporal lobe function during memory encoding. The Journal of Neuroscience: The Official Journal of the Society for Neuroscience, 30(28), 9548–9556. https://doi.org/10.1523/JNEUROSCI.3500-09.2010

Ghetti, S., & Fandakova, Y. (2020). Neural Development of Memory and Metamemory in Childhood and Adolescence: Toward an Integrative Model of the Development of Episodic Recollection. Annual Review of Developmental Psychology, 2(1), 365–388. https://doi.org/10.1146/annurev-devpsych-060320-085634

Gonsalves, B. D., Kahn, I., Curran, T., Norman, K. A., & Wagner, A. D. (2005). Memory strength and repetition suppression: multimodal imaging of medial temporal cortical contributions to recognition. Neuron, 47(5), 751–761. https://doi.org/10.1016/j.neuron.2005.07.013

Hallquist, M. N., Geier, C. F., & Luna, B. (2018). Incentives facilitate developmental improvement in inhibitory control by modulating control-related networks. NeuroImage, 172, 369–380. https://doi.org/10.1016/j.neuroimage.2018.01.045

Hanten, G., Li, X., Chapman, S. B., Swank, P., Gamino, J., Roberson, G., & Levin, H. S. (2007). Development of verbal selective learning. Developmental Neuropsychology, 32(1), 585–596. https://doi.org/10.1080/87565640701361112

Hennessee, J. P., Castel, A. D., & Knowlton, B. J. (2017). Recognizing What Matters: Value Improves Recognition by Selectively Enhancing Recollection. Journal of Memory and Language, 94, 195–205. https://doi.org/10.1016/j.jml.2016.12.004

Hennessee, J. P., Patterson, T. K., Castel, A. D., & Knowlton, B. J. (2019). Forget me not: Encoding processes in value-directed remembering. In Journal of Memory and Language (Vol. 106, pp. 29–39). https://doi.org/10.1016/j.jml.2019.02.001

Hofer, S. M., Flaherty, B. P., & Hoffman, L. (2006). Cross-sectional analysis of time-dependent data: Mean-induced association in age-heterogeneous samples and an alternative method based on sequential narrow age-cohort samples. Multivariate Behavioral Research, 41(2), 165–187. https://doi.org/10.1207/s15327906mbr4102_4

Insel, C., Kastman, E. K., Glenn, C. R., & Somerville, L. H. (2017). Development of corticostriatal connectivity constrains goal-directed behavior during adolescence. Nature Communications, 8(1), 1605. https://doi.org/10.1038/s41467-017-01369-8

Jang, A. I., Nassar, M. R., Dillon, D. G., & Frank, M. J. (2019). Positive reward prediction errors during decision-making strengthen memory encoding. Nature Human Behaviour, 3(7), 719–732. https://doi.org/10.1038/s41562-019-0597-3

Jenkinson, M., Beckmann, C. F., Behrens, T. E. J., Woolrich, M. W., & Smith, S. M. (2012). FSL. NeuroImage, 62(2), 782–790. https://doi.org/10.1016/j.neuroimage.2011.09.015

Jonides, J., & Naveh-Benjamin, M. (1987). Estimating frequency of occurrence. Journal of Experimental Psychology. Learning, Memory, and Cognition, 13(2), 230–240. https://doi.org/10.1037/0278-7393.13.2.230

Katzman, P. L., & Hartley, C. A. (2020). The value of choice facilitates subsequent memory across development. Cognition, 199, 104239. https://doi.org/10.1016/j.cognition.2020.104239

Kirchhoff, B. A., & Buckner, R. L. (2006). Functional-anatomic correlates of individual differences in memory. Neuron, 51(2), 263–274. https://doi.org/10.1016/j.neuron.2006.06.006

Kirchhoff, B. A., Wagner, A. D., Maril, A., & Stern, C. E. (2000). Prefrontal–Temporal Circuitry for Episodic Encoding and Subsequent Memory. The Journal of Neuroscience: The Official Journal of the Society for Neuroscience, 20(16), 6173–6180. https://doi.org/10.1523/JNEUROSCI.20-16-06173.2000

Kleiner, M., Brainard, D., & Pelli, D. (2007). What’s new in Psychtoolbox-3? https://pure.mpg.de/rest/items/item_1790332/component/file_3136265/content

Köhler, S., Danckert, S., Gati, J. S., & Menon, R. S. (2005). Novelty responses to relational and non-relational information in the hippocampus and the parahippocampal region: A comparison based on event-related fMRI. In Hippocampus (Vol. 15, Issue 6, pp. 763y8774). https://doi.org/10.1002/hipo.20098

Kundu, P., Brenowitz, N. D., Voon, V., Worbe, Y., Vértes, P. E., Inati, S. J., Saad, Z. S., Bandettini, P. A., & Bullmore, E. T. (2013). Integrated strategy for improving functional connectivity mapping using multiecho fMRI. Proceedings of the National Academy of Sciences of the United States of America, 110(40), 16187–16192. https://doi.org/10.1073/pnas.1301725110

Kundu, P., Inati, S. J., Evans, J. W., Luh, W.-M., & Bandettini, P. A. (2012). Differentiating BOLD and non-BOLD signals in fMRI time series using multi-echo EPI. NeuroImage, 60(3), 1759–1770. https://doi.org/10.1016/j.neuroimage.2011.12.028

Libby, R., & Lipe, M. G. (1992). Incentives, Effort, and the Cognitive Processes Involved in Accounting-Related Judgments. Journal of Accounting Research, 30(2), 249–273. https://doi.org/10.2307/2491126

Liljeholm, M., & O’Doherty, J. P. (2012). Contributions of the striatum to learning, motivation, and performance: an associative account. Trends in Cognitive Sciences, 16(9), 467–475. https://doi.org/10.1016/j.tics.2012.07.007

Lindenberger, U., von Oertzen, T., Ghisletta, P., & Hertzog, C. (2011). Cross-sectional age variance extraction: what’s change got to do with it? Psychology and Aging, 26(1), 34–47. https://doi.org/10.1037/a0020525

Lisman, J. E., & Grace, A. A. (2005). The hippocampal-VTA loop: controlling the entry of information into long-term memory. Neuron, 46(5), 703–713. https://doi.org/10.1016/j.neuron.2005.05.002

Liu, Y., Mattar, M. G., Behrens, T. E. J., Daw, N. D., & Dolan, R. J. (2020). Experience replay supports non-local learning. In Cold Spring Harbor Laboratory (p. 2020.10.20.343061). https://doi.org/10.1101/2020.10.20.343061

Mathworks Inc. (2017). MATLAB: R2017a. Mathworks Inc, Natick.

Meulemans, T., Van der Linden, M., & Perruchet, P. (1998). Implicit sequence learning in children. Journal of Experimental Child Psychology, 69(3), 199–221. https://doi.org/10.1006/jecp.1998.2442

Miotto, E. C., Savage, C. R., Evans, J. J., Wilson, B. A., Martins, M. G. M., Iaki, S., & Amaro, E., Jr. (2006). Bilateral activation of the prefrontal cortex after strategic semantic cognitive training. Human Brain Mapping, 27(4), 288–295. https://doi.org/10.1002/hbm.20184

Moeller, S., Yacoub, E., Olman, C. A., Auerbach, E., Strupp, J., Harel, N., & Uğurbil, K. (2010). Multiband multislice GE-EPI at 7 tesla, with 16-fold acceleration using partial parallel imaging with application to high spatial and temporal whole-brain fMRI. In Magnetic Resonance in Medicine (Vol. 63, Issue 5, pp. 1144–1153). https://doi.org/10.1002/mrm.22361

Murty, V. P., & Adcock, R. A. (2014). Enriched encoding: reward motivation organizes cortical networks for hippocampal detection of unexpected events. Cerebral Cortex, 24(8), 2160–2168. https://doi.org/10.1093/cercor/bht063

Murty, V. P., Calabro, F., & Luna, B. (2016). The role of experience in adolescent cognitive development: Integration of executive, memory, and mesolimbic systems. Neuroscience and Biobehavioral Reviews, 70, 46–58. https://doi.org/10.1016/j.neubiorev.2016.07.034

Murty, V. P., Tompary, A., Adcock, R. A., & Davachi, L. (2017). Selectivity in Postencoding Connectivity with High-Level Visual Cortex Is Associated with Reward-Motivated Memory. The Journal of Neuroscience: The Official Journal of the Society for Neuroscience, 37(3), 537–545. https://doi.org/10.1523/JNEUROSCI.4032-15.2016

Ngo, C., Newcombe, N., & Olson, I. R. (2019). Gain-loss framing enhances mnemonic discrimination in preschoolers. Child Development, 90(5), 1569–1578. https://doi.org/10.31234/osf.io/nxq9t

Nordt, M., Hoehl, S., & Weigelt, S. (2016). The use of repetition suppression paradigms in developmental cognitive neuroscience. Cortex; a Journal Devoted to the Study of the Nervous System and Behavior, 80, 61–75. https://doi.org/10.1016/j.cortex.2016.04.002

Nussenbaum, K., Prentis, E., & Hartley, C. A. (2020). Memory’s reflection of learned information value increases across development. Journal of Experimental Psychology: General. https://psycnet.apa.org/record/2020-18143-001

Ofen, N. (2012). The development of neural correlates for memory formation. Neuroscience and Biobehavioral Reviews, 36(7), 1708–1717. https://doi.org/10.1016/j.neubiorev.2012.02.016

Ofen, N., Kao, Y.-C., Sokol-Hessner, P., Kim, H., Whitfield-Gabrieli, S., & Gabrieli, J. D. E. (2007). Development of the declarative memory system in the human brain. Nature Neuroscience, 10(9), 1198–1205. https://doi.org/10.1038/nn1950

O’Kane, G., Insler, R. Z., & Wagner, A. D. (2005). Conceptual and perceptual novelty effects in human medial temporal cortex. Hippocampus, 15(3), 326–332. https://doi.org/10.1002/hipo.20053

Pachur, T., Schooler, L. J., & Stevens, J. R. (2014). We’ll meet again: revealing distributional and temporal patterns of social contact. PloS One, 9(1), e86081. https://doi.org/10.1371/journal.pone.0086081

Pelli, D. G. (1997). The VideoToolbox software for visual psychophysics: transforming numbers into movies. Spatial Vision, 10(4), 437–442. https://www.ncbi.nlm.nih.gov/pubmed/9176953

Popov, V., & Reder, L. M. (2020). Frequency effects on memory: A resource-limited theory. Psychological Review, 127(1), 1–46. https://doi.org/10.1037/rev0000161

Power, J. D., & Petersen, S. E. (2013). Control-related systems in the human brain. Current Opinion in Neurobiology, 23(2), 223–228. https://doi.org/10.1016/j.conb.2012.12.009

R Core Team. (2018). R: A language and environment for statistical computing (Version 3.5.1) [Computer software]. https://www.R-project.org

Reder, L. M., Liu, X. L., Keinath, A., & Popov, V. (2016). Building knowledge requires bricks, not sand: The critical role of familiar constituents in learning. Psychonomic Bulletin & Review, 23(1), 271–277. https://doi.org/10.3758/s13423-015-0889-1

Rich, A. S., & Gureckis, T. M. (2018). Exploratory choice reflects the future value of information. Decisions, 5(3), 177–192. https://doi.org/10.1037/dec0000074

Rosenbaum, G., Grassie, H., & Hartley, C. A. (2020). Valence biases in reinforcement learning shift across adolescence and modulate subsequent memory. https://doi.org/10.31234/osf.io/n3vsr

Rouhani, N., Norman, K. A., & Niv, Y. (2018). Dissociable effects of surprising rewards on learning and memory. Journal of Experimental Psychology. Learning, Memory, and Cognition, 44(9), 1430–1443. https://doi.org/10.1037/xlm0000518

Scherf, K. S., Luna, B., Avidan, G., & Behrmann, M. (2011). “What” Precedes “Which”: Developmental Neural Tuning in Face- and Place-Related Cortex. Cerebral Cortex, 21(9), 1963–1980. https://doi.org/10.1093/cercor/bhq269

Scimeca, J. M., & Badre, D. (2012). Striatal contributions to declarative memory retrieval. Neuron, 75(3), 380–392. https://doi.org/10.1016/j.neuron.2012.07.014

Shadlen, M. N., & Shohamy, D. (2016). Decision Making and Sequential Sampling from Memory. Neuron, 90(5), 927–939. https://doi.org/10.1016/j.neuron.2016.04.036

Shigemune, Y., Tsukiura, T., Kambara, T., & Kawashima, R. (2014). Remembering with gains and losses: effects of monetary reward and punishment on successful encoding activation of source memories. Cerebral Cortex, 24(5), 1319–1331. https://doi.org/10.1093/cercor/bhs415

Shing, Y. L., Brehmer, Y., Heekeren, H. R., Bäckman, L., & Lindenberger, U. (2016). Neural activation patterns of successful episodic encoding: Reorganization during childhood, maintenance in old age. Developmental Cognitive Neuroscience, 20, 59–69. https://doi.org/10.1016/j.dcn.2016.06.003

Shing, Y. L., Werkle-Bergner, M., Brehmer, Y., Müller, V., Li, S.-C., & Lindenberger, U. (2010). Episodic memory across the lifespan: the contributions of associative and strategic components. Neuroscience and Biobehavioral Reviews, 34(7), 1080–1091. https://doi.org/10.1016/j.neubiorev.2009.11.002

Shohamy, D., & Adcock, R. A. (2010). Dopamine and adaptive memory. Trends in Cognitive Sciences, 14(10), 464–472. https://doi.org/10.1016/j.tics.2010.08.002

Simonsohn, U. (2018). Two lines: A valid alternative to the invalid testing of U-shaped relationships with quadratic regressions. Advances in Methods and Practices in Psychological Science, 1(4), 538–555. https://doi.org/10.1177/2515245918805755

Singmann, H., Bolker, B., Westfall, J., Aust, F., & Ben-Shachar, M. S. (2020). Afex: analysis of factorial experiments (Version. 27-2) [Computer software]. https://CRAN.R-project.org/package=afex

Smith, S. M., Jenkinson, M., Woolrich, M. W., Beckmann, C. F., Behrens, T. E. J., Johansen-Berg, H., Bannister, P. R., De Luca, M., Drobnjak, I., Flitney, D. E., Niazy, R. K., Saunders, J., Vickers, J., Zhang, Y., De Stefano, N., Brady, J. M., & Matthews, P. M. (2004). Advances in functional and structural MR image analysis and implementation as FSL. NeuroImage, 23 Suppl 1, S208–19. https://doi.org/10.1016/j.neuroimage.2004.07.051

Somerville, L. H., & Casey, B. J. (2010). Developmental neurobiology of cognitive control and motivational systems. Current Opinion in Neurobiology, 20(2), 236–241. https://doi.org/10.1016/j.conb.2010.01.006

Somerville, L. H., Jones, R. M., Ruberry, E. J., Dyke, J. P., Glover, G., & Casey, B. J. (2013). The medial prefrontal cortex and the emergence of self-conscious emotion in adolescence. Psychological Science, 24(8), 1554–1562. https://doi.org/10.1177/0956797613475633

Squire, L. R. (1992). Declarative and nondeclarative memory: multiple brain systems supporting learning and memory. Journal of Cognitive Neuroscience, 4(3), 232–243. https://doi.org/10.1162/jocn.1992.4.3.232

Stanek, J. K., Dickerson, K. C., Chiew, K. S., Clement, N. J., & Adcock, R. A. (2019). Expected Reward Value and Reward Uncertainty Have Temporally Dissociable Effects on Memory Formation. Journal of Cognitive Neuroscience, 31(10), 1443–1454. https://doi.org/10.1162/jocn_a_01411

Stevens, J. R., Marewski, J. N., Schooler, L. J., & Gilby, I. C. (2016). Reflections of the social environment in chimpanzee memory: applying rational analysis beyond humans. Royal Society Open Science, 3(8), 160293. https://doi.org/10.1098/rsos.160293

Stokes, M. G., Atherton, K., Patai, E. Z., & Nobre, A. C. (2012). Long-term memory prepares neural activity for perception. Proceedings of the National Academy of Sciences of the United States of America, 109(6), E360–7. https://doi.org/10.1073/pnas.1108555108

Störmer, V., Eppinger, B., & Li, S.-C. (2014). Reward speeds up and increases consistency of visual selective attention: a lifespan comparison. Cognitive, Affective & Behavioral Neuroscience, 14(2), 659–671. https://doi.org/10.3758/s13415-014-0273-z

Summerfield, J. J., Lepsien, J., Gitelman, D. R., Mesulam, M. M., & Nobre, A. C. (2006). Orienting attention based on long-term memory experience. Neuron, 49(6), 905–916. https://doi.org/10.1016/j.neuron.2006.01.021

Tang, L., Shafer, A. T., & Ofen, N. (2018). Prefrontal Cortex Contributions to the Development of Memory Formation. Cerebral Cortex, 28(9), 3295–3308. https://doi.org/10.1093/cercor/bhx200

Tingley, D., Yamamoto, T., Hirose, K., Keele, L., & Imai, K. (2014). Mediation: R package for causal mediation analysis. Journal of Statistical Software. https://oar.princeton.edu/jspui/handle/88435/pr1gj2f

Turi, M., Burr, D. C., Igliozzi, R., Aagten-Murphy, D., Muratori, F., & Pellicano, E. (2015). Children with autism spectrum disorder show reduced adaptation to number. Proceedings of the National Academy of Sciences of the United States of America, 112(25), 7868–7872. https://doi.org/10.1073/pnas.1504099112

Turk-Browne, N. B., Yi, D.-J., & Chun, M. M. (2006). Linking implicit and explicit memory: common encoding factors and shared representations. Neuron, 49(6), 917–927. https://doi.org/10.1016/j.neuron.2006.01.030

Uncapher, M. R., Hutchinson, J. B., & Wagner, A. D. (2011). Dissociable effects of top-down and bottom-up attention during episodic encoding. The Journal of Neuroscience: The Official Journal of the Society for Neuroscience, 31(35), 12613–12628. https://doi.org/10.1523/JNEUROSCI.0152-11.2011

Uncapher, M. R., & Wagner, A. D. (2009). Posterior parietal cortex and episodic encoding: insights from fMRI subsequent memory effects and dual-attention theory. Neurobiology of Learning and Memory, 91(2), 139–154. https://doi.org/10.1016/j.nlm.2008.10.011

Wais, P. E., Kim, O. Y., & Gazzaley, A. (2012). Distractibility during episodic retrieval is exacerbated by perturbation of left ventrolateral prefrontal cortex. Cerebral Cortex, 22(3), 717–724. https://doi.org/10.1093/cercor/bhr160

Wang, S., Feng, S. F., & Bornstein, A. (2020). Mixing memory and desire: How memory reactivation supports deliberative decision-making. PsyArXiv. https://doi.org/10.31234/osf.io/5vksj

Ward, E. J., Chun, M. M., & Kuhl, B. A. (2013). Repetition suppression and multi-voxel pattern similarity differentially track implicit and explicit visual memory. The Journal of Neuroscience: The Official Journal of the Society for Neuroscience, 33(37), 14749–14757. https://doi.org/10.1523/JNEUROSCI.4889-12.2013

Wechsler, D. (2011). Wechsler Abbreviated Scale of Intelligence (WASI-II). NCS Pearson.

Wittmann, B. C., Schott, B. H., Guderian, S., Frey, J. U., Heinze, H.-J., & Düzel, E. (2005). Reward-related FMRI activation of dopaminergic midbrain is associated with enhanced hippocampus-dependent long-term memory formation. Neuron, 45(3), 459–467. https://doi.org/10.1016/j.neuron.2005.01.010

Woolrich, M. W., Ripley, B. D., Brady, M., & Smith, S. M. (2001). Temporal autocorrelation in univariate linear modeling of FMRI data. NeuroImage, 14(6), 1370–1386. https://doi.org/10.1006/nimg.2001.0931

Woolrich, Mark W., Behrens, T. E. J., Beckmann, C. F., Jenkinson, M., & Smith, S. M. (2004). Multilevel linear modelling for FMRI group analysis using Bayesian inference. NeuroImage, 21(4), 1732–1747. https://doi.org/10.1016/j.neuroimage.2003.12.023

Xu, J., Moeller, S., Auerbach, E. J., Strupp, J., Smith, S. M., Feinberg, D. A., Yacoub, E., & Uğurbil, K. (2013). Evaluation of slice accelerations using multiband echo planar imaging at 3 T. NeuroImage, 83, 991–1001. https://doi.org/10.1016/j.neuroimage.2013.07.055

Yu, Q., McCall, D. M., Homayouni, R., Tang, L., Chen, Z., Schoff, D., Nishimura, M., Raz, S., & Ofen, N. (2018). Age-associated increase in mnemonic strategy use is linked to prefrontal cortex development. Neuroimage, 181, 162–169. https://doi.org/10.1016/j.neuroimage.2018.07.008

