## Supplemental materials for "Developmental change in prefrontal cortex recruitment supports the emergence of value-guided memory"

### Supplementary Participant Information

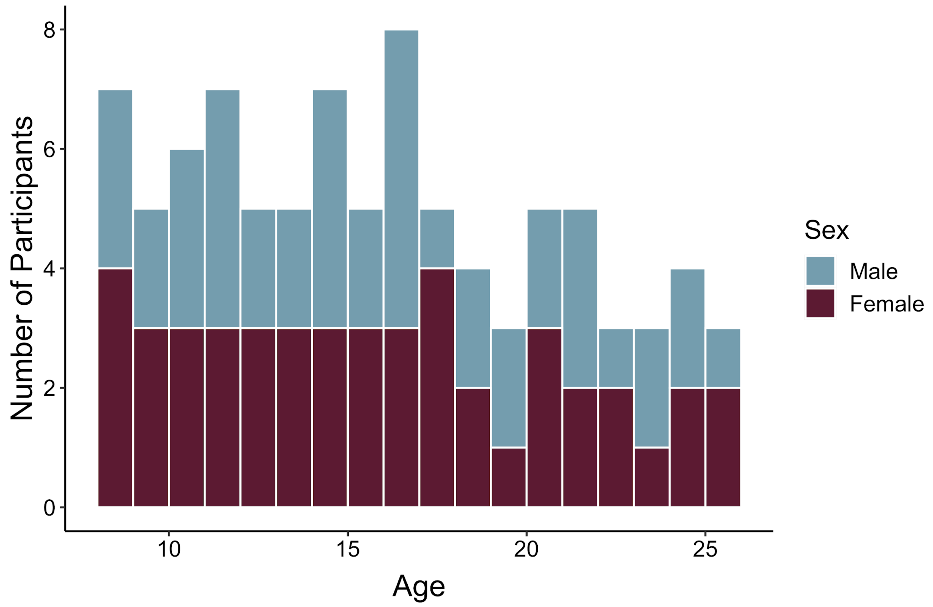


**Figure S1.** Participant age and sex distribution.

**Table S1**: Number of participants included in each analysis

| Block | Data Type | Frequency-learning | Associative encoding | Retrieval | Frequency Reports |
|---|---|---|---|---|---|
| 1 | Behavioral | 89 | *NA* | 90 | 90 |
| 1 | Neural | 88 | 90 | 90 | *NA* |
| 2 | Behavioral | 86 | *NA* | 85 | 85 |
| 2 | Neural | 84 | 81 | 81 | *NA* |

### Supplementary Results

#### Frequency-learning: Accuracy and reaction times

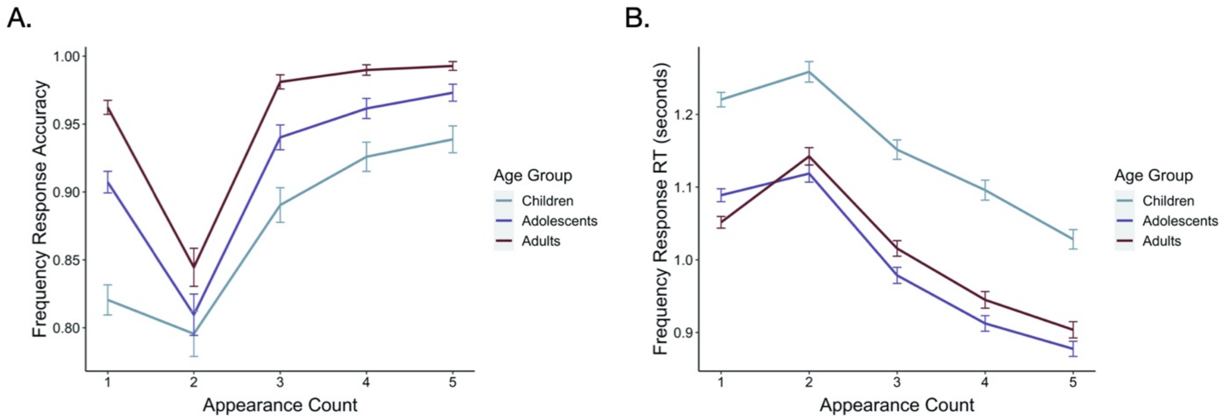


**Figure S2.** (A) During frequency-learning, older participants were more accurate in identifying items as new ($\chi2(1) = 25.54$, p <.001) and as repeated ($\chi2(1) = 33.81$, p < .001). All participants became more accurate in identifying items as repeated as the number of repetitions increased ($\chi2 = 138.20$, *p* < .001), though younger participants demonstrated a greater increase in accuracy throughout learning ($\chi2(1) = 17.52$, p < .001). (B) Older participants also responded to both new ($F(1, 85.99) = 32.51$, $p < .001$) and repeated ($F(1, 87.55) = 21.82$, $p < .001$) items more quickly than younger participants. Reaction times to old items became faster as the a function of item repetition number ($F(1, 69.94) = 282.21$, $p < .001$).

#### Relation between age and associative memory: Two-lines test

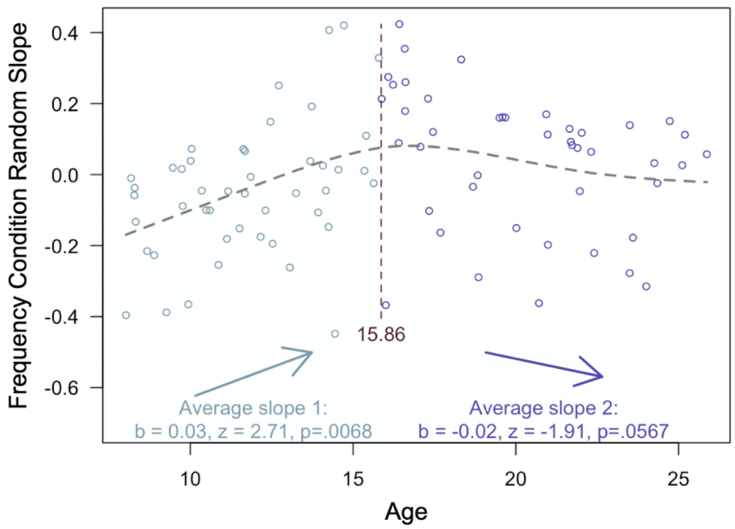


**Figure S3.** Results from the two-lines test (Simonsohn, 2018) revealed that the influence of frequency condition on memory accuracy increased throughout childhood and early adolescence, and did not significantly decrease from adolescence into early adulthood.

### Prefrontal cortex activation during encoding

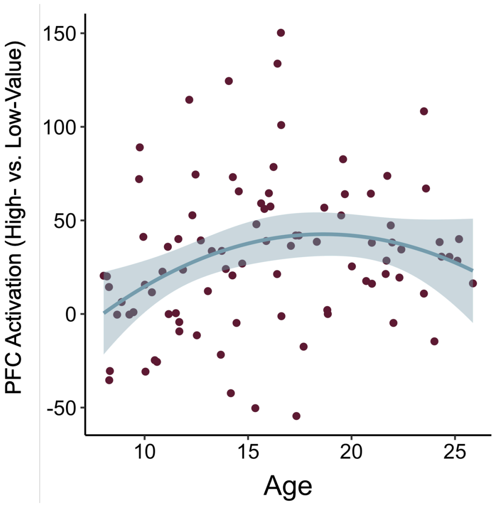


**Figure S4.** Mean beta weights averaged over voxels within a prefrontal cortex ROI (see 'methods' in main text) during encoding of associations involving high- vs. low-frequency items increased with age. The increase was greatest in childhood before leveling out into late adolescence and early adulthood.

### Hippocampal and parahippocampal cortex activation during encoding

*A priori,* we expected that regions in the medial temporal lobe that have been linked to successful memory formation, including the hippocampus and parahippocampal cortex (Davachi, 2006), may be differentially engaged during encoding of high- vs. low- value information. Further, we hypothesized that the differential engagement of these regions across age may contribute to age differences in value-guided memory. Though we did not see any significant clusters of activation in the hippocampus or parahippocampal cortex in our group level high value vs. low value encoding contrast, we conducted additional ROI analyses to test these hypotheses. As with our other ROI analyses, we first identified the peak voxel (based on its $z$-statistic; hippocampus: $x = 24$, $y = 34$, $z = 23$; parahippocampal cortex: $x = 22$, $y = 41$, $z = 16$) in each region from our group-level contrast, and then drew 5-mm spheres around them. We then examined how average parameter estimates within these spheres related to both age and memory difference scores.

First, we ran a linear regression modeling the effects of age, WASI scores, and their interaction on hippocampal activation. We did not observe a main effect of age on hippocampal activation, ($\beta = .00$, SE = .10, $p > .99$). We did, however, observe a significant age x WASI score interaction effect ($\beta = .30$, SE = .10, $p = .003$). Next, we conducted another linear regression to examine the effects of hippocampal activation, age, WASI scores, and their interaction on memory difference scores. In contrast to our prefrontal cortex activation results, activation in the hippocampus did not relate to memory difference scores, ($\beta = -.02$, SE = .03, $p = .50$).

We repeated these analyses with our parahippocampal cortex sphere. Here, we did not observe any significant effects of age on parahippocampal activation ($\beta = -.07$, SE = .11, $p = .50$), nor did we observe any effects of parahippocampal activation on memory difference scores ($\beta = .01$, SE = .03, $p = .25$).

**Effects of block order and type on associative memory**

*Block order.* To examine whether participants' memory, or their use of learned value to guide memory, varied across blocks, we re-ran our associative memory accuracy model with block order (e.g., 1 or 2) as an additional interacting fixed effect. Our full model included frequency condition, WASI score, linear and quadratic age, and block order as interacting fixed effects. We included random intercepts and random slopes across frequency condition and block order for each participant, and random intercepts and random slopes across frequency condition, IQ, linear and quadratic age, and block order for each stimulus. We did not observe a significant effect of block order on associative memory ($p$ = .676; Table S2), nor did block order interact with any other predictors ($p$s > .18). Thus, we did not observe any evidence that participants performed the task differently across blocks.

*Block type.* To examine whether participants' memory, or their use of learned value to guide memory, varied across blocks depending on their content, we re-ran our associative memory accuracy model with block type (e.g., pictures/frames or postcards/stamps) as an additional interacting fixed effect. Our full model included frequency condition, WASI score, linear and quadratic age, and block type as interacting fixed effects. We included random intercepts and random slopes across frequency condition and block type for each participant, and random intercepts and random slopes across frequency condition, IQ, linear and quadratic age, and block type for each stimulus. We did not observe a main effect of block type on associative memory ($p$ = .061; Table S3; Figure S5). We did, however, observe a significant block type x frequency condition interaction, such that participants better remembered low-value pairs in the block with the pictures. Importantly, when we included block type as a covariate, we continued to observe a robust influence of frequency condition on associative memory, as well as significant interactions between frequency condition and both age terms.

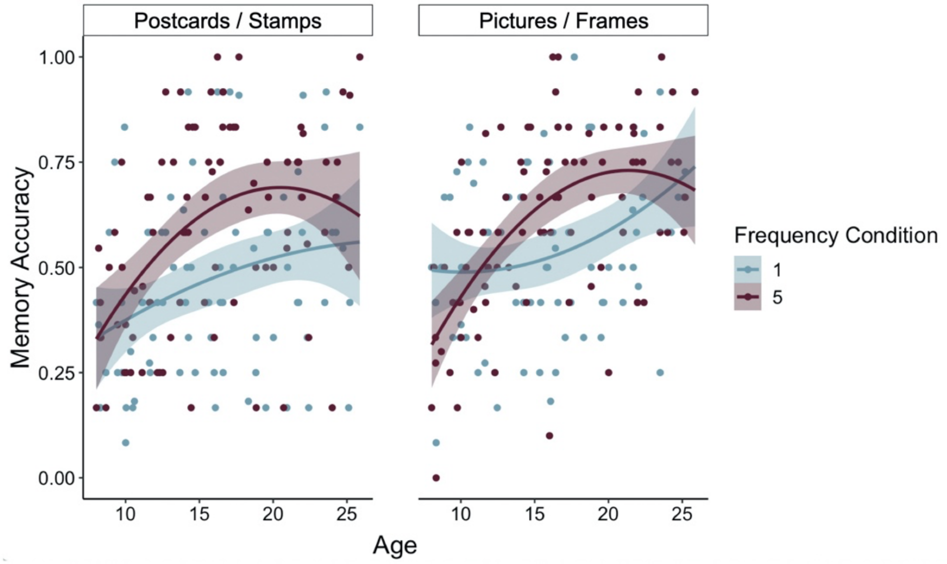


**Figure S5**. Participants demonstrated a greater influence of frequency condition on associative memory in the task block involving postcards and stamps relative to the task block involving pictures and frames ($\chi2(1)$ =.4.40, p = .036).

**Table S2:** Associative memory accuracy by frequency condition with block order

| | ***Estimate*** | ***95% CI*** | $X^2$ | ***p*** |
|---|---|---|---|---|
| Intercept | 0.26 | 0.12 – 0.40 | | |
| **Age** | **1.42** | **0.52 -2.33** | **8.97** | **.003** |
| **Age$^2$** | **-0.97** | **-1.87 - -0.08** | **4.39** | **.036** |
| **WASI** | **0.28** | **0.14 – 0.41** | **15.09** | **<.001** |
| **Frequency Condition** | **-0.22** | **-0.31 - -0.13** | **19.47** | **<.001** |
| Block Order | -0.02 | -0.11 – 0.07 | 0.18 | .676 |
| Age x WASI | 0.15 | -0.80 – 1.10 | 0.10 | .756 |
| Age$^2$ x WASI | -0.10 | -1.00 – 0.80 | 0.04 | .833 |
| Age x Block Order | -0.23 | -0.85 – 0.39 | 0.53 | .468 |
| Age$^2$ x Block Order | 0.13 | -0.48 – 0.75 | 0.18 | .675 |
| **Age x Frequency Condition** | **-1.05** | **-1.68 - -0.43** | **10.31** | **.001** |
| **Age$^2$ x Frequency Condition** | **0.97** | **-0.35 – 1.59** | **8.94** | **.003** |
| WASI x Frequency Condition | -0.04 | -0.13 – 0.05 | 0.65 | .419 |
| Block Order x Frequency Condition | -0.03 | -0.10 – 0.05 | 0.76 | .384 |
| Block Order x WASI | -0.05 | -0.15 – 0.04 | 1.13 | .288 |
| Age x WASI x Frequency Condition | -0.53 | -1.19 – 0.13 | 2.43 | .119 |
| Age$^2$ x WASI x Frequency Condition | 0.54 | -0.08 – 1.17 | 2.83 | .092 |
| Age x Block Order x Frequency Condition | -0.35 | -0.86 – 0.16 | 1.80 | .180 |
| Age$^2$ x Block Order x Frequency Condition | 0.27 | -0.23 – 0.77 | 1.10 | .294 |
| WASI x Block Order x Frequency Condition | -0.05 | -0.12 – 0.03 | 1.35 | .246 |
| WASI x Age x Block Order | 0.05 | -0.61 – 0.70 | 0.02 | .887 |
| WASI x Age$^2$ x Block Order | -0.01 | -0.63 – 0.62 | 0.00 | .985 |
| Age x WASI x Frequency Condition x Block Order | 0.35 | -0.19 – 0.89 | 1.61 | .205 |
| Age$^2$ x WASI x Frequency Condition x Block Order | -0.33 | -0.85 – 0.18 | 1.62 | .203 |

**Table S3:** Associative memory accuracy by frequency condition with block type

| | ***Estimate*** | ***95% CI*** | $X^2$ | ***p*** |
|---|---|---|---|---|
| Intercept | 0.26 | 0.12 – 0.40 | | |
| **Age** | **1.40** | **0.49 – 2.31** | **8.61** | **.003** |
| **Age$^2$** | **-0.95** | **-1.85 - -0.05** | **4.16** | **.041** |
| **WASI** | **0.27** | **0.14 – 0.40** | **14.46** | **<.001** |
| **Frequency Condition** | **-0.22** | **-0.31 - -0.13** | **20.07** | **<.001** |
| Block Type | -0.10 | -0.20 – 0.00 | 3.52 | .061 |
| Age x WASI | 0.17 | -0.79 – 1.12 | 0.12 | .734 |
| Age$^2$ x WASI | -0.10 | -1.01 – 0.80 | 0.05 | .822 |
| Age x Block Type | 0.23 | -0.39 – 0.85 | 0.53 | .465 |
| Age$^2$ x Block Type | -0.28 | -0.89 – 0.33 | 0.79 | .375 |
| **Age x Frequency Condition** | **-1.05** | **-1.68 - -0.42** | **10.06** | **.002** |
| **Age$^2$ x Frequency Condition** | **0.96** | **0.34 – 1.59** | **8.64** | **.003** |
| WASI x Frequency Condition | -0.04 | -0.13 – 0.05 | 0.65 | .419 |
| **Block Type x Frequency Condition** | **-0.08** | **-0.15 - -0.01** | **4.40** | **.036** |
| Block Type x WASI | 0.01 | -0.08 – 0.10 | 0.03 | .866 |
| Age x WASI x Frequency Condition | -0.51 | -1.18 – 0.15 | 2.24 | .135 |
| Age$^2$ x WASI x Frequency Condition | 0.52 | -0.11 – 1.15 | 2.56 | .109 |
| Age x Block Type x Frequency Condition | 0.31 | -0.19 – 0.82 | 1.44 | .230 |
| Age$^2$ x Block Type x Frequency Condition | -0.29 | -0.79 – 0.21 | 1.28 | .258 |
| WASI x Block Type x Frequency Condition | -0.03 | -0.10 – 0.05 | 0.44 | .505 |
| WASI x Age x Block Type | 0.62 | -0.03 – 1.27 | 3.44 | .064 |
| WASI x Age$^2$ x Block Type | -0.60 | -1.22 – 0.02 | 3.57 | .059 |
| Age x WASI x Frequency Condition x Block Type | -0.31 | -0.84 – 0.23 | 1.28 | .258 |
| Age$^2$ x WASI x Frequency Condition x Block Type | 0.34 | -0.17 – 0.85 | 1.68 | .195 |

Given that we observed a significant block type x frequency condition interaction on memory, we next examined whether the relation between age and lateral PFC activation during encoding varied across block type. To do so, we used fslmeants to extract the mean parameter estimate within our PFC ROI for the 5 vs. 1 encoding contrast for each participant, for each block. We then examined how these parameter estimates varied with age. To do so, we ran a linear mixed-effects model with linear age, quadratic age, WASI scores, and block type as interacting fixed effects, and included random participant intercepts. We did not observe a significant effect of block type on PFC activation, nor did it interact with any other predictors (*ps* > .22). In line with the analyses reported in the main text, we observed a significant relation between linear age and PFC activation ($p = .007$; Table S4).

**Table S4**: High- vs. low-value encoding PFC activation by age with block type

| | *Estimate* | *95% CI* | *df* | *F* | *p* |
|---|---|---|---|---|---|
| Intercept | -.01 | -0.17 - 0.15 | | | |
| **Age** | **1.60** | **0.47 – 2.73** | **1, 79.46** | **7.76** | **.007** |
| **Age$^2$** | **-1.39** | **-2.50 - -0.28** | **1, 79.24** | **6.04** | **.016** |
| WASI | 0.15 | -0.02 – 0.32 | 1, 86.82 | 3.06 | .084 |
| Block Type | -0.04 | -0.19 – 0.11 | 1, 80.30 | 0.30 | .585 |
| Age x WASI | 0.45 | -0.79 – 1.69 | 1, 87.35 | 0.50 | .479 |
| Age$^2$ x WASI | -0.53 | -1.69 – 0.63 | 1, 86.03 | 0.79 | .376 |
| Age x Block Type | 0.11 | -0.94 – 1.16 | 1, 78.81 | 0.04 | .843 |
| Age$^2$ x Block Type | -0.12 | -1.16 – 0.91 | 1, 78.59 | 0.05 | .818 |
| WASI x Block Type | -0.06 | -0.22 – 0.10 | 1, 86.37 | 0.54 | .462 |
| Age x WASI x Block Type | -0.73 | -1.89 – 0.44 | 1, 86.58 | 1.49 | .226 |
| Age$^2$ x WASI x Block Type | 0.68 | -0.41 – 1.77 | 1, 85.26 | 1.48 | .227 |

Finally, we examined how PFC activation across blocks influenced memory difference scores via a linear mixed-effects model with PFC activation, age, WASI scores, and block type as interacting fixed effects. Our model also included random participant intercepts. Here, including quadratic age did not improve model fit ($X^2(8) = 0.00, p = 1$). We found that PFC activation related to memory difference scores ($p = .002$; Table S5). No other main effects or interactions were significant (*p*s > .063).

**Table S5**: Memory difference scores by PFC activation and age with block type

| | *Estimate* | *95% CI* | *df* | *F* | *p* |
|---|---|---|---|---|---|
| Intercept | 0.09 | 0.05 – 0.13 | | | |
| **PFC Activation** | **0.06** | **0.02 – 0.10** | **1, 153.78** | **9.85** | **.002** |
| Age | 0.02 | -0.01 – 0.06 | 1, 84.35 | 1.52 | .221 |
| WASI | -0.00 | -0.04 – 0.04 | 1, 86.86 | 0.04 | .836 |
| Block Type | 0.03 | -0.00 – 0.07 | 1, 82.48 | 2.92 | .091 |
| PFC Activation x Age | 0.01 | -0.03 – 0.06 | 1, 153.53 | 0.28 | .598 |
| PFC Activation x WASI | 0.03 | -0.01 – 0.08 | 1, 154.29 | 2.13 | .147 |
| Age x WASI | -0.01 | -0.05 – 0.03 | 1, 86.37 | 0.22 | .642 |
| PFC Activation x Block Type | -0.01 | -0.04 – 0.03 | 1, 151.41 | 0.09 | .770 |
| Age x Block Type | 0.01 | -0.03 – 0.04 | 1, 82.91 | 0.13 | .718 |
| WASI x Block Type | 0.02 | -0.02 – 0.05 | 1, 85.40 | 0.76 | .385 |
| PFC Activation x Age x WASI | -0.01 | -0.06 – 0.04 | 1, 152.59 | 0.14 | .711 |
| PFC Activation x Age x Block Type | 0.03 | -0.02 – 0.08 | 1, 152.38 | 1.56 | .214 |
| PFC Activation x WASI x Block Type | -0.04 | -0.09 – 0.00 | 1, 151.64 | 3.49 | .064 |
| Age x WASI x Block Type | -0.02 | -0.05 – 0.01 | 1, 85.20 | 1.23 | .270 |
| PFC Activation x Age x WASI x Block Type | 0.05 | -0.01 – 0.10 | 1, 154.25 | 3.00 | .085 |

**Full Model Specification and Results**

For each model described in the manuscript, we report here its full random-effects structure (when relevant) and effect estimates.

**Model 1: Frequency-learning accuracy: new items.** We examined how participants' accuracy in identifying new items during frequency-learning varied as a function of age, WASI scores, and their interaction via a mixed-effects logistic regression. We included random intercepts for each participant and each stimulus. Including quadratic age in the model did not improve model fit ($X^2(2) = 3.29$, $p = .19$).

**Table S6:** Frequency-learning accuracy: new items

| | *Estimate* | *95% CI* | $X^2$ | *p* |
|---|---|---|---|---|
| Intercept | 3.08 | 2.73 - 3.44 | | |
| **Age** | **0.92** | **0.58 – 1.25** | **25.52** | **<.001** |
| **WASI** | **0.43** | **0.09 – 0.77** | **6.18** | **.013** |
| Age x WASI | 0.25 | -0.08 – 0.57 | 2.25 | .134 |

**Model 2**: **Frequency-learning accuracy: repeated items.** We examined how participants' accuracy in identifying repeated items during frequency-learning varied as a function of the number of times the item had appeared, age, WASI scores, and their interactions via a mixed-effects logistic regression. We included random intercepts and random slopes across item appearances for each participant and random intercepts for each stimulus. Including quadratic age did not improve model fit ($X^2(4) = 1.80$, $p = .77$).

**Table S7:** Frequency-learning accuracy: repeated item appearances

| | *Estimate* | *95% CI* | $X^2$ | *p* |
|---|---|---|---|---|
| Intercept | 3.83 | 3.46 – 4.20 | | |
| **Appearance** | **1.53** | **1.28 – 1.78** | **138.03** | **<.001** |
| **Age** | **0.97** | **0.64 – 1.29** | **33.43** | **<.001** |
| **WASI** | **0.46** | **0.13 – 0.79** | **7.58** | **.006** |
| **Appearance x Age** | **0.45** | **0.24 – 0.67** | **17.41** | **<.001** |
| Appearance x WASI | 0.05 | -0.17 – 0.26 | 0.18 | .672 |
| Age x WASI | 0.12 | -0.20 – 0.45 | 0.57 | .449 |
| Appearance x Age x WASI | 0.04 | -0.17 – 0.25 | 0.14 | .707 |

**Model 3: Frequency-learning reaction times: new items**. We examined how participants' reaction times when they correctly identified new items during frequency-learning varied as a function of age, WASI scores, and their interaction via a mixed-effects linear regression. We included random intercepts for each participant and each stimulus, and random slopes across age and WASI scores for each stimulus. We also estimated the correlation between random stimulus intercepts and slopes. Including quadratic age did not improve model fit ($X^2(13) = 21.26, p = .068$).

**Table S8:** Frequency-learning reaction times: new items

| | *Estimate* | *95% CI* | *df* | *F* | *p* |
|---|---|---|---|---|---|
| Intercept | 1.12 | 1.09 - 1.15 | | | |
| **Age** | **-0.08** | **-0.11 - -0.05** | **1, 85.99** | **32.51** | **<.001** |
| WASI | -0.01 | -0.04 - 0.02 | 1, 82.34 | 0.56 | .457 |
| Age x WASI | -0.02 | -0.05 - -0.01 | 1, 83.14 | 2.12 | .149 |

**Model 4: Frequency-learning reaction times: repeated items.** We examined how participants' reaction times when they correctly identified repeated items during frequency-learning varied as a function of the number of times the item had appeared, age, WASI scores, and their interactions via a mixed-effects linear regression. We included random intercepts and random slopes across item appearances for each participant, random intercepts and slopes across age, WASI scores, and item appearances for each stimulus, and estimated the correlation between random stimulus intercepts and slopes. Including quadratic age did not improve model fit ($X^2(9) = 3.18, p = .96$).

**Table S9:** Frequency-learning reaction times: repeated items

| | *Estimate* | *95% CI* | *df* | *F* | *p* |
|---|---|---|---|---|---|
| Intercept | 1.03 | 1.00 – 1.06 | | | |
| **Age** | **-0.07** | **-0.10 – -0.04** | **1, 87.55** | **21.82** | **<.001** |
| WASI | -0.03 | -0.06 – -0.01 | 1, 86.27 | 2.65 | .108 |
| **Appearance** | **-0.08** | **-0.09 – -0.07** | **1, 69.94** | **282.21** | **<.001** |
| Age x WASI | -0.01 | -0.03 – 0.02 | 1, 84.97 | 0.22 | .641 |
| Age x Appearance | -0.01 | -0.01 – 0.00 | 1, 77.06 | 1.26 | .265 |
| WASI x Appearance | 0.00 | -0.01 – 0.01 | 1, 75.79 | 0.00 | .992 |
| Age x WASI x Appearance | 0.00 | -0.01 – 0.01 | 1, 74.96 | 0.68 | .413 |

**Model 5: Parahippocampal cortex neural activation by stimulus repetition and age.** For items in the high-frequency condition, we examined how neural activation in a parahippocampal cortex ROI varied as a function of age, quadratic age, stimulus repetition number, quadratic stimulus repetition number, WASI scores, and their interactions. We included random intercepts for each participant and stimulus, random slopes across linear and quadratic repetition number for each participant, and random slopes across linear and quadratic repetition number, linear and quadratic age, and their interactions for each stimulus stimuli.

**Table S10:** Parahippocampal cortex neural activation by stimulus repetition and age

| | ***Estimate*** | ***95% CI*** | ***df*** | ***F*** | ***p*** |
|---|---|---|---|---|---|
| Intercept | 65.70 | 52.23 – 79.17 | | | |
| Age | -78.41 | -164.25 – 7.43 | 1, 82.78 | 3.21 | .077 |
| Age$^2$ | 82.54 | -2.27 – 167.35 | 1, 82.77 | 3.64 | .060 |
| **Repetition** | **-30.20** | **-40.89 – -19.50** | **1, 5015.94** | **30.64** | **<.001** |
| **Repetition$^2$** | **14.52** | **4.10 – 24.93** | **1, 9881.00** | **7.47** | **.006** |
| WASI | -1.11 | -13.49 – 11.27 | 1, 83.31 | 0.03 | .861 |
| **Age x Repetition** | **101.65** | **27.40 – 175.90** | **1, 7267.46** | **7.20** | **.007** |
| **Age x Repetition$^2$** | **-88.18** | **161.49 - -14.87** | **1, 9857.85** | **5.56** | **.018** |
| **Age$^2$ x Repetition** | **97.99** | **-171.28 - -24.71** | **1, 7260.70** | **6.87** | **.009** |
| **Age$^2$ x Repetition$^2$** | **82.76** | **10.40 – 155.11** | **1, 9854.92** | **5.03** | **.025** |
| WASI x Age | 28.87 | -61.72 – 119.47 | 1, 83.51 | 0.39 | .534 |
| WASI x Age$^2$ | -20.73 | -106.34 – 64.88 | 1, 83.47 | 0.23 | .636 |
| WASI x Repetition | -7.56 | -18.46 – 3.35 | 1, 7402.99 | 1.84 | .175 |
| WASI x Repetition$^2$ | 7.40 | -3.38 – 18.18 | 1, 7857.10 | 1.81 | .178 |
| WASI x Age x Repetition | -52.32 | -130.26 – 25.61 | 1, 7243.45 | 1.73 | .188 |
| WASI x Age x Repetition$^2$ | 42.15 | -34.79 – 119.08 | 1, 9868.65 | 1.15 | .283 |
| WASI x Age$^2$ x Repetition | 45.97 | -27.58 – 119.53 | 1, 7235.30 | 1.50 | .221 |
| WASI x Age$^2$ x Repetition$^2$ | -38.01 | -110.62 – 34.59 | 1, 9867.59 | 1.05 | .305 |

**Model 6: Repetition suppression indices and age.** For items in the high-frequency condition, we examined how repetition suppression varied as a function of age, quadratic age, WASI scores, and their interactions. We included random intercepts for each participant and stimulus, and random slopes across age, quadratic age, and WASI scores for each stimulus.

**Table S11:** Repetition suppression indices

| | *Estimate* | *95% CI* | *df* | *F* | *p* |
|---|---|---|---|---|---|
| Intercept | 45.54 | 35.63 – 55.45 | | | |
| **Age** | **-61.34** | **-8.56 – 10.77** | **1, 78.32** | **3.97** | **.050** |
| **Age$^2$** | **66.52** | **-121.70 - -0.98** | **1, 77.55** | **4.80** | **.031** |
| WASI | 1.11 | 7.01 – 126.03 | 1, 58.06 | 0.05 | .823 |
| Age x WASI | 60.90 | -2.53 – 124.34 | 1, 77.38 | 3.54 | .064 |
| Age$^2$ x WASI | -51.17 | -111.03 – 8.70 | 1, 77.16 | 2.81 | .098 |

**Model 7: Frequency report error magnitudes.** We examined how the magnitude of participants' errors in their frequency reports varied as a function of age, WASI scores, frequency condition, and their interactions via a mixed-effects linear regression. We included random intercepts and random slopes across frequency conditions for each participant and random intercepts and random slopes across age, WASI scores, and frequency conditions for each stimulus. Including quadratic age did not improve model fit ($\chi^2(8) = 7.96, p = .437$).

**Table S12:** Frequency report error magnitudes

| | *Estimate* | *95% CI* | *df* | *F* | *p* |
|---|---|---|---|---|---|
| Intercept | 1.21 | 1.12 – 1.30 | | | |
| **Age** | **-0.18** | **-0.27 - - 0.10** | **1, 94.30** | **17.57** | **<.001** |
| **WASI** | **-0.11** | **-0.19 - -0.02** | **1, 83.80** | **6.47** | **.014** |
| Frequency Condition | -0.00 | -0.11 – 0.11 | 1, 93.81 | 0.00 | .993 |
| Age x WASI | -0.07 | -0.15 – 0.01 | 1, 86.35 | 3.24 | .075 |
| Age x Frequency Condition | -0.05 | -0.17 – 0.06 | 1, 85.48 | 0.95 | .332 |
| WASI x Frequency Condition | -0.03 | -0.14 – 0.09 | 1, 86.55 | 0.20 | .652 |
| Age x WASI x Frequency Condition | -0.07 | -0.17 – 0.03 | 1, 85.94 | 1.77 | .187 |

**Model 8: Frequency reports by repetition suppression indices.** For items in the high-frequency condition, we examined how frequency reports varied as a function of age, WASI scores, repetition suppression, and their interactions via a mixed-effects linear regression. We included random intercepts and random slopes across repetition suppression for each participant, and random intercepts and random slopes across repetition suppression, WASI scores, age, and their interactions for each stimulus. Including a quadratic age term did not improve model fit ($\chi^2(8)$ = 6.45, $p$ = .60).

**Table S13:** Frequency reports by repetition suppression

| | *Estimate* | *95% CI* | *df* | *F* | *p* |
|---|---|---|---|---|---|
| Intercept | 4.44 | 4.26 - 4.63 | | | |
| **Age** | **0.26** | **0.09 - 0.42** | **1, 82.87** | **8.93** | **.004** |
| **WASI** | **0.19** | **0.02 - 0.36** | **1, 85.19** | **4.77** | **.032** |
| Repetition Suppression | 0.00 | -0.06 - 0.07 | 1, 1360.74 | 0.01 | .903 |
| Age x WASI | 0.09 | -0.07 - 0.25 | 1, 84.13 | 1.34 | .251 |
| Age x Repetition Suppression | 0.06 | -0.00 - 0.12 | 1, 938.87 | 3.61 | .058 |
| WASI x Repetition Suppression | 0.04 | -0.02 - 0.10 | 1, 58.81 | 1.52 | .222 |
| Age x WASI x Repetition Suppression | -0.03 | -0.08 - 0.03 | 1, 313.72 | 1.10 | .296 |

**Model 9: Associative memory accuracy**. We examined how memory accuracy varied as a function of age, quadratic age, WASI scores, frequency condition, and their interactions via a mixed-effects logistic regression. We included random intercepts and random slopes across frequency conditions for each participant, and random intercepts and random slopes across frequency condition, WASI scores, age, and quadratic age for each stimulus.

**Table S14:** Associative memory accuracy by frequency condition

| | ***Estimate*** | ***95% CI*** | $\chi^2$ | ***p*** |
|---|---|---|---|---|
| Intercept | 0.26 | 0.12 – 0.40 | | |
| **Age** | **1.38** | **0.49 – 2.28** | **8.68** | **.003** |
| **Age$^2$** | **-0.95** | **-1.83 – -0.06** | **4.24** | **.039** |
| **WASI** | **0.26** | **0.13 – 0.39** | **14.18** | **<.001** |
| **Frequency Condition** | **-0.21** | **-0.30 – -0.13** | **19.73** | **<.001** |
| Age x WASI | 0.18 | -0.76 – 1.12 | 0.14 | .704 |
| Age$^2$ x WASI | -0.12 | -1.01 – 0.77 | 0.07 | .789 |
| **Age x Frequency Condition** | **-1.06** | **-1.68 – -0.45** | **10.74** | **.001** |
| **Age$^2$ x Frequency Condition** | **0.98** | **0.37 – 1.59** | **9.27** | **.002** |
| WASI x Frequency Condition | -0.04 | -0.13 – 0.05 | 0.86 | .355 |
| Age x WASI x Frequency Condition | -0.50 | -1.15 – 0.15 | 2.26 | .133 |
| Age$^2$ x WASI x Frequency Condition | 0.52 | -0.10 – 1.13 | 2.65 | .104 |

**Model 10: Associative memory accuracy excluding participants who performed below chance.** Two participants (both children) responded correctly to 25% or fewer memory test trials. We re-ran our memory accuracy model, excluding these two participants.

**Table S15:** Associative memory accuracy by frequency condition (below-chance subjects excluded)

| | *Estimate* | *95% CI* | $X^2$ | *p* |
|---|---|---|---|---|
| Intercept | 0.30 | 0.16 – 0.44 | | |
| **Age** | **1.19** | **0.29 – 2.09** | **6.48** | **.011** |
| Age$^2$ | -0.79 | -1.68 – 0.10 | 2.98 | .084 |
| **WASI** | **0.24** | **0.11 – 0.37** | **12.44** | **<.001** |
| **Frequency Condition** | **-0.22** | **-0.31 – -0.13** | **20.04** | **<.001** |
| Age x WASI | 0.31 | -0.62 - 1.24 | 0.43 | .513 |
| Age$^2$ x WASI | -0.23 | -1.11 - 0.66 | 0.25 | .615 |
| **Age x Frequency Condition** | **-1.07** | **-1.70 - -0.43** | **10.25** | **.001** |
| **Age$^2$ x Frequency Condition** | **0.99** | **0.36 - 1.61** | **8.97** | **.003** |
| WASI x Frequency Condition | -0.04 | -0.14 - 0.05 | 0.81 | .368 |
| Age x WASI x Frequency Condition | -0.50 | -1.16 - 0.16 | 2.17 | .141 |
| Age$^2$ x WASI x Frequency Condition | 0.52 | -0.11 - 1.15 | 2.54 | .111 |

**Model 11: Associative memory accuracy controlling for individual differences in frequency learning.** We examined how memory accuracy varied as a function of age, quadratic age, WASI scores, frequency condition, and their interactions via a mixed-effects logistic regression. We also included mean frequency report error magnitudes, overall mean accuracy during frequency-learning, and mean accuracy on the last appearance of each item during frequency-learning as non-interacting fixed effects. We included random intercepts and random slopes across frequency conditions for each participant, and random intercepts and random slopes across frequency condition, WASI scores, age, and quadratic age for each stimulus.

**Table S16:** Associative memory accuracy by frequency condition (with frequency-learning covariates)

| | *Estimate* | *95% CI* | $X^2$ | *p* |
|---|---|---|---|---|
| Intercept | 0.25 | 0.12 – 0.38 | | |
| Age | 0.59 | -0.31 – 1.49 | 1.62 | .203 |
| Age$^2$ | -0.30 | -1.17 – 0.56 | 0.47 | .491 |
| **WASI** | **0.16** | **0.03 – 0.29** | **6.02** | **.014** |
| **Frequency Condition** | **-0.21** | **-0.30 – -0.13** | **19.65** | **<.001** |
| **Mean Frequency Report Error Magnitude** | **-0.26** | **-0.39 – -0.12** | **13.05** | **<.001** |
| Frequency-learning Accuracy | 0.17 | -0.04 – 0.38 | 2.36 | .125 |
| Frequency-learning Accuracy (last item appearance) | -0.10 | -0.30 – 0.09 | 1.06 | .304 |
| Age x WASI | 0.11 | -0.75 – 0.96 | 0.06 | .807 |
| Age$^2$ x WASI | -0.08 | -0.89 – 0.72 | 0.04 | .843 |
| **Age x Frequency Condition** | **-1.06** | **-1.67 - -0.44** | **10.59** | **.001** |
| **Age$^2$ x Frequency Condition** | **0.97** | **0.36 - 1.58** | **9.15** | **.002** |
| WASI x Frequency Condition | -0.04 | -0.13 - 0.05 | 0.83 | .362 |
| Age x WASI x Frequency Condition | -0.50 | -1.15 - 0.15 | 2.22 | .136 |
| Age$^2$ x WASI x Frequency Condition | 0.51 | -0.10 - 1.13 | 2.61 | .106 |

**Model 12: High- vs. low-value encoding caudate activation and age.** We ran a linear regression to examine the effects of age, WASI scores, and their interaction on differential caudate activation during encoding of high- vs. low-value information. Including quadratic age did not improve model fit ($X^2(2) = 2.26$, $p = .11$).

**Table S17:** High vs. low-value encoding caudate activation by age

| | *Estimate* | *SE* | *t* | *p* |
|---|---|---|---|---|
| Intercept | -0.07 | .104 | | |
| Age | 0.16 | .107 | 1.55 | .126 |
| WASI | 0.18 | .110 | 1.64 | .105 |
| **Age x WASI** | **-0.29** | **.101** | **-2.86** | **.005** |

**Model 13: High- vs. low-value encoding PFC activation and age.** We ran a linear regression to examine the effects of linear and quadratic age, WASI scores, and their interactions on differential PFC activation during encoding of high- vs. low-value information.

**Table S18:** High vs. low-value encoding PFC activation by age

| | *Estimate* | *SE* | *t* | *p* |
|---|---|---|---|---|
| Intercept | 0.00 | .105 | | |
| **Age** | **1.97** | **.743** | **2.65** | **.009** |
| **Age$^2$** | **-1.73** | **.734** | **-2.35** | **.021** |
| **WASI** | **0.26** | **.109** | **2.34** | **.022** |
| Age x WASI | 0.93 | .789 | 1.18 | .240 |
| Age$^2$ x WASI | -1.02 | .745 | -1.37 | .174 |

**Model 14: Relation between frequency reports and associative memory accuracy.** We examined how memory accuracy varied as a function of age, quadratic age, WASI scores, frequency reports, and their interactions via a mixed-effects logistic regression. We included random intercepts and random slopes across frequency reports for each participant, and random intercepts and random slopes across frequency reports, age, quadratic age, and WASI scores for each stimulus.

**Table S19:** Associative memory accuracy by frequency report

| | ***Estimate*** | ***95% CI*** | $X^2$ | ***p*** |
|---|---|---|---|---|
| Intercept | 0.26 | 0.12 – 0.40 | | |
| **Age** | **1.46** | **0.55 – 2.38** | **9.25** | **.002** |
| **Age$^2$** | **-1.01** | **-1.92 – -0.11** | **4.64** | **.031** |
| **WASI** | **0.27** | **0.13 – 0.40** | **13.94** | **<.001** |
| **Frequency Report** | **0.28** | **0.19 – 0.37** | **31.20** | **<.001** |
| Age x WASI | 0.15 | -0.81 – 1.12 | 0.09 | .759 |
| Age$^2$ x WASI | -0.10 | -1.01 – 0.82 | 0.04 | .838 |
| **Age x Frequency Report** | **1.13** | **0.46 – 1.79** | **10.37** | **.001** |
| **Age$^2$ x Frequency Report** | **-1.07** | **-1.73 – -0.41** | **9.50** | **.002** |
| WASI x Frequency Report | 0.02 | -0.08 – 0.11 | 0.10 | .754 |
| Age x WASI x Frequency Report | 0.37 | -0.35 – 1.09 | 1.00 | .316 |
| Age$^2$ x WASI x Frequency Report | -0.33 | -1.01 – 0.35 | 0.89 | .345 |

**Model 15: Influence of repetition suppression on associative memory accuracy.** For associations involving items in the high-frequency condition, we examined how memory accuracy varied as a function of age, quadratic age, WASI scores, repetition suppression, and their interactions via a mixed-effects logistic regression. We included random intercepts and random slopes across repetition suppression for each participant, and random intercepts and random slopes across repetition suppression, WASI scores, age, quadratic age, and their interactions for each stimulus.

**Table S20:** Associative memory accuracy by repetition suppression

| | ***Estimate*** | ***95% CI*** | $\mathcal{X}^2$ | ***p*** |
|---|---|---|---|---|
| Intercept | 0.51 | 0.32 – 0.69 | | |
| **Age** | **2.63** | **1.43 – 3.83** | **16.87** | **<.001** |
| **Age$^2$** | **-2.13** | **-3.31 – -0.94** | **11.55** | **<.001** |
| **WASI** | **0.31** | **0.14 – 0.49** | **11.47** | **<.001** |
| **Repetition Suppression** | **0.23** | **0.10 – 0.37** | **11.21** | **<.001** |
| Age x WASI | 0.78 | -0.49 – 2.05 | 1.44 | .230 |
| Age$^2$ x WASI | -0.75 | -1.95 – 0.45 | 1.47 | .225 |
| Age x Repetition Suppression | -0.42 | -1.32 – 0.49 | 0.79 | .374 |
| Age$^2$ x Repetition Suppression | 0.64 | -0.28 – 1.57 | 1.79 | .181 |
| WASI x Repetition Suppression | 0.00 | -0.13 – 0.14 | 0.00 | .954 |
| Age x WASI x Repetition Suppression | -0.17 | -1.02 – 0.68 | 0.15 | .700 |
| Age$^2$ x WASI x Repetition Suppression | 0.27 | -0.57 – 1.10 | 0.37 | .541 |

**Model 16: Effects of frequency reports and repetition suppression on associative memory accuracy.** For associations involving items in the high-frequency condition, we examined how memory accuracy varied as a function of age, quadratic age, WASI scores, repetition suppression, frequency reports, and their interactions via a mixed-effects logistic regression. We included random intercepts and random slopes across repetition suppression, frequency reports, and their interaction for each participant. We also included random intercepts and random slopes across repetition suppression, frequency reports, age, quadratic age, and WASI scores for each stimulus.

**Table S21:** Associative memory accuracy by repetition suppression and frequency reports

| | *Estimate* | *95% CI* | $X^2$ | *p* |
|---|---|---|---|---|
| Intercept | 0.51 | 0.32 – 0.69 | | |
| **Repetition Suppression** | **0.23** | **0.09 – 0.36** | **10.25** | **.001** |
| **WASI** | **0.26** | **0.08 – 0.44** | **7.59** | **.006** |
| **Frequency Report** | **0.3** | **0.17 – 0.42** | **21.16** | **<.001** |
| **Age** | **2.47** | **1.25 – 3.69** | **14.4** | **<.001** |
| **Age$^2$** | **-2.02** | **-3.23 – -0.81** | **9.98** | **.002** |
| Repetition Suppression x WASI | 0.00 | -0.13 – 0.14 | 0.00 | .968 |
| Repetition Suppression x Frequency Report | -0.11 | -0.25 – 0.04 | 2.18 | .140 |
| WASI x Frequency Report | -0.05 | -0.18 – 0.08 | 0.48 | .488 |
| Repetition Suppression x Age | -0.12 | -1.05 – 0.81 | 0.06 | .804 |
| Repetition Suppression x Age$^2$ | 0.33 | -0.62 – 1.27 | 0.45 | .503 |
| WASI x Age | 0.92 | -0.36 – 2.21 | 1.96 | .161 |
| WASI x Age$^2$ | -0.89 | -2.12 – 0.33 | 2.04 | .153 |
| Frequency Report x Age | 0.12 | -0.74 – 0.99 | 0.08 | .783 |
| Frequency Report x Age$^2$ | -0.09 | -0.97 – 0.79 | 0.04 | .843 |
| RS x WASI x Frequency Report | -0.04 | -0.19 – 0.11 | 0.27 | .603 |
| RS x WASI x Age | -0.28 | -1.16 – 0.61 | 0.36 | .550 |
| RS x WASI x Age$^2$ | 0.39 | -0.49 – 1.26 | 0.71 | .400 |
| RS x Frequency Report x Age | 0.13 | -0.78 – 1.03 | 0.07 | .786 |
| RS x Frequency Report x Age$^2$ | -0.12 | -1.09 – 0.85 | 0.06 | .809 |
| WASI x Frequency Report x Age | -0.33 | -1.31 – 0.65 | 0.44 | .509 |
| WASI x Frequency Report x Age$^2$ | 0.44 | -0.52 – 1.40 | 0.79 | .374 |
| RS x WASI x Frequency Report x Age | -0.73 | -1.68 – 0.22 | 2.29 | .130 |
| RS x WASI x Frequency Report x Age$^2$ | 0.83 | -0.14 – 1.80 | 2.86 | .091 |

**Model 17: Relation between age and mean repetition suppression indices.** We ran a linear regression to examine how average repetition suppression indices (across items, for each participant) varied as a function of age, quadratic age, WASI scores, and their interactions.

**Table S22.** Mean repetition suppression indices by age

| | ***Estimate*** | ***SE*** | ***t*** | ***p*** |
|---|---|---|---|---|
| Intercept | 44.76 | 4.53 | | |
| Age | -59.00 | 31.82 | -1.85 | .067 |
| **Age$^2$** | **64.93** | **31.37** | **2.07** | **.042** |
| WASI | 1.86 | 4.72 | 0.39 | .695 |
| Age x WASI | 57.06 | 33.89 | 1.68 | .096 |
| Age$^2$ x WASI | -48.02 | 31.93 | -1.50 | .136 |

**Model 18: Relation between mean repetition suppression indices and neural activation in caudate.** We ran a linear regression to examine how differential caudate activation in response to high- vs. low-value information during encoding related to average repetition suppression indices age, WASI scores, and their interactions. Including quadratic age did not improve model fit ($X^2(4) = 1.37$, $p = .25$).

**Table S23:** Caudate activation by repetition suppression indices

| | ***Estimate*** | ***SE*** | ***t*** | ***p*** |
|---|---|---|---|---|
| Intercept | 8.23 | 1.54 | | |
| Repetition Suppression | 2.31 | 1.60 | 1.44 | .153 |
| Age | 1.48 | 1.64 | 0.91 | .367 |
| WASI | 1.93 | 1.64 | 1.17 | .244 |
| Repetition Suppression x Age | 1.29 | 1.42 | 0.91 | .366 |
| Repetition Suppression x WASI | 0.26 | 1.54 | 0.17 | .864 |
| **Age x WASI** | **-4.40** | **1.50** | **-2.94** | **.004** |
| Repetition Suppression x Age x WASI | -0.09 | 1.22 | -0.07 | .945 |

**Model 19: Relation between mean repetition suppression indices and PFC neural activation.** We ran a linear regression to examine how differential PFC activation in response to high- vs. low-value information during encoding related to average repetition suppression indices, age, quadratic age, WASI scores, and their interactions.

**Table S24:** PFC activation by repetition suppression indices

| | ***Estimate*** | ***SE*** | ***t*** | ***p*** |
|---|---|---|---|---|
| Intercept | 28.66 | 4.51 | | |
| Repetition Suppression | -6.78 | 4.83 | -1.40 | .165 |
| **Age** | **103.36** | **35.33** | **2.93** | **.004** |
| **Age$^2$** | **-94.25** | **35.13** | **-2.68** | **.009** |
| **WASI** | **13.59** | **4.74** | **2.87** | **.005** |
| Repetition Suppression x Age | -54.15 | 34.13 | -1.59 | .112 |
| Repetition Suppression x Age$^2$ | 53.85 | 32.79 | 1.64 | .105 |
| Repetition Suppression x WASI | -3.28 | 4.49 | -0.73 | .467 |
| Age x WASI | 30.43 | 34.44 | 0.88 | .380 |
| Age$^2$ x WASI | -38.59 | 32.41 | -1.91 | .237 |
| Repetition Suppression x Age x WASI | -41.33 | 29.80 | -1.39 | .169 |
| Repetition Suppression x Age$^2$ x WASI | 41.46 | 27.58 | 1.50 | .137 |

**Model 20: Relation between age and frequency distance.** We ran a linear regression to examine how mean frequency distances related to age, quadratic age, WASI scores, and their interactions.

**Table S25.** Frequency distance by age

| | *Estimate* | *SE* | *T* | *p* |
|---|---|---|---|---|
| Intercept | 2.23 | .09 | | |
| **Age** | **2.75** | **.67** | **4.10** | **<.001** |
| **Age$^2$** | **-2.28** | **.66** | **-3.44** | **<.001** |
| **WASI** | **0.38** | **.10** | **3.89** | **<.001** |
| Age x WASI | 0.33 | .71 | 0.46 | .646 |
| Age$^2$ x WASI | -0.12 | .67 | -0.18 | .857 |

**Model 21: Relation between frequency distance and neural activation in caudate.** We ran a linear regression to examine how differential caudate activation in response to high- vs. low-value information during encoding related to average repetition suppression indices age, WASI scores, and their interactions. Including quadratic age did not improve model fit ($X^2(4) = 1.36$, $p = .25$).

**Table S26.** Caudate activation by frequency distance

| | *Estimate* | *SE* | *t* | *p* |
|---|---|---|---|---|
| Intercept | 7.00 | 1.78 | | |
| Frequency Distance | 2.55 | 1.86 | 1.37 | .175 |
| Age | 1.39 | 1.89 | 0.74 | .463 |
| WASI | 1.13 | 1.77 | 0.64 | .526 |
| Frequency Distance x Age | 2.27 | 1.77 | 1.29 | .202 |
| Frequency Distance x WASI | -1.50 | 1.77 | -0.85 | .399 |
| **Age x WASI** | **-5.49** | **1.64** | **-3.43** | **.001** |
| Frequency Distance x Age x WASI | 1.71 | 1.40 | 1.22 | .227 |

**Model 22: Relation between frequency distance and PFC neural activation**. We ran a linear regression to examine how differential PFC activation in response to high- vs. low-value information during encoding related to frequency distance, age, WASI scores, and their interactions. Including quadratic age did not improve model fit ($X^2(4) = 1.49$, $p = .21$).

**Table S27.** PFC activation by frequency distance

| | *Estimate* | *SE* | *t* | *p* |
|---|---|---|---|---|
| Intercept | -0.02 | .12 | | |
| **Frequency Distance** | **0.42** | **.12** | **3.36** | **.001** |
| Age | -0.03 | .13 | -0.22 | .824 |
| WASI | 0.08 | .12 | 0.64 | .522 |
| Frequency Distance x Age | -0.18 | .12 | -1.51 | .136 |
| Frequency Distance x WASI | 0.15 | .12 | 1.23 | .223 |
| Age x WASI | -0.17 | .11 | -1.52 | .132 |
| Frequency Distance x Age x WASI | 0.05 | .09 | 0.52 | .607 |

**Model 23: Effects of frequency distance and PFC neural activation on associative memory accuracy.** We ran a linear regression to examine how associative memory accuracy was related to differential PFC activation in response to high- vs. low-value information during encoding, frequency distance, age, quadratic age, WASI scores, and their interactions.

**Table S28.** Associative memory accuracy by PFC activation and frequency distance

| | ***Estimate*** | ***SE*** | ***t*** | ***p*** |
|---|---|---|---|---|
| Intercept | 0.07 | | | |
| Frequency Distance | -0.02 | 0.03 | -0.55 | .582 |
| **Age** | **0.56** | **0.2** | **2.83** | **.006** |
| **Age$^2$** | **-0.51** | **0.2** | **-2.6** | **.012** |
| **WASI** | 0.02 | 0.03 | 0.85 | .398 |
| **PFC Activation** | 0.08 | 0.04 | 1.95 | .055 |
| Frequency Distance x Age | 0.29 | 0.2 | 1.5 | .139 |
| Frequency Distance x Age$^2$ | -0.27 | 0.2 | -1.36 | .178 |
| Frequency Distance x WASI | 0.01 | 0.03 | 0.38 | .709 |
| Age x WASI | 0.03 | 0.19 | 0.17 | .866 |
| Age$^2$ x WASI | -0.05 | 0.18 | -0.25 | .800 |
| Frequency Distance x PFC Activation | 0.01 | 0.04 | 0.25 | .800 |
| Age x PFC Activation | -0.15 | 0.34 | -0.43 | .672 |
| Age$^2$ x PFC Activation | 0.21 | 0.34 | 0.62 | .538 |
| WASI x PFC Activation | -0.04 | 0.04 | -0.84 | .406 |
| Frequency Distance x Age x WASI | -0.36 | 0.22 | -1.66 | .102 |
| Frequency Distance x Age$^2$ x WASI | 0.33 | 0.2 | 1.62 | .111 |
| Frequency Distance x Age x PFC Activation | -0.07 | 0.27 | -0.24 | .809 |
| Frequency Distance x Age$^2$ x PFC Activation | 0.08 | 0.28 | 0.28 | .778 |
| Frequency Distance x WASI x PFC Activation | -0.05 | 0.03 | -1.49 | .142 |
| Age$^2$ x WASI x PFC Activation | 0.46 | 0.36 | 1.29 | .202 |
| Age$^2$ x WASI x PFC Activation | -0.46 | 0.36 | -1.28 | .204 |
| Frequency Distance x Age$^2$ x WASI x PFC Activation | 0.27 | 0.27 | 0.99 | .324 |
| Frequency Distance x Age$^2$ x WASI x PFC Activation | -0.28 | 0.27 | -1.07 | .288 |

### Supplemental Neural Results

**Table S29:** Frequency-learning: Last vs. first item appearance cluster table

| ***Region*** | ***x*** | ***y*** | ***z*** | ***Cluster size*** | ***z-max*** |
|---|---|---|---|---|---|
| Frontal pole | -42 | 51 | 0 | 2419 | 6.33 |
| Precuneus | 0 | -66 | 33 | 1322 | 8.99 |
| Left lateral occipital cortex / angular gyrus | -57 | -66 | 30 | 1319 | 7.2 |
| Right lateral occipital cortex / angular gyrus | 51 | -63 | 33 | 637 | 6.03 |
| Right middle temporal gyrus | 66 | -33 | -12 | 304 | 5.48 |
| Right cerebellum | 15 | -87 | -27 | 164 | 5.33 |
| Precentral gyrus | 3 | -18 | 75 | 124 | 4.53 |
| Left cerebellum | -42 | -75 | -42 | 92 | 4.72 |
| Left middle temporal gyrus | -57 | -3 | -27 | 73 | 4.44 |
| Left caudate | -6 | 12 | 9 | 62 | 4.91 |
| Right caudate | 9 | 24 | 6 | 60 | 5.04 |
| Occipital pole | -3 | -96 | 12 | 46 | 4.81 |

**Table S30:** Frequency-learning: First vs. last item appearance cluster table

| ***Region*** | ***x*** | ***y*** | ***z*** | ***Cluster size*** | ***z-max*** |
|---|---|---|---|---|---|
| Right temporal fusiform cortex / lateral occipital cortex / parahippocampal gyrus | 30 | -39 | -15 | 2137 | 8.62 |
| Left temporal fusiform cortex / lateral occipital cortex / parahippocampal gyrus | -33 | -63 | -15 | 1858 | 7.53 |
| Cingulate gyrus | 9 | 9 | 42 | 100 | 6 |
| Right precuneus | 18 | -51 | 9 | 80 | 6.05 |
| Left postcentral gyrus | -51 | -15 | 57 | 74 | 4.73 |
| Right precentral gyrus | 42 | 3 | 33 | 73 | 5.54 |
| Juxtapositional lobule cortex | 6 | 6 | 57 | 24 | 4.11 |
| Left amygdala | -24 | -6 | -15 | 22 | 4.23 |
| Cingulate gyrus | 3 | -3 | 33 | 22 | 4.68 |
| Central opercular cortex | 39 | -3 | 15 | 21 | 4.8 |

**Table S31:** Encoding: Encoding vs. baseline by linear age cluster table

| ***Region*** | ***x*** | ***y*** | ***z*** | ***Cluster size*** | ***z-max*** |
|---|---|---|---|---|---|
| Right lateral occipital cortex | 45 | -81 | 21 | 132 | 5 |
| Left precentral gyrus / middle frontal gyrus | -54 | 12 | 33 | 120 | 5.9 |
| Left lateral occipital cortex | -45 | -84 | 21 | 54 | 4.69 |
| Left lateral occipital cortex | -27 | -72 | 42 | 50 | 4.32 |
| Right lateral occipital cortex | 30 | -69 | 45 | 44 | 4.17 |
| Left cerebellum | -18 | -45 | -48 | 39 | 4.55 |
| Superior frontal gyrus | -3 | 12 | 60 | 39 | 4.35 |
| Left supramarginal gyrus | -36 | -45 | 36 | 37 | 4.57 |
| Left middle frontal gyrus | -33 | 3 | 63 | 33 | 4.6 |
| Right superior parietal lobule | 33 | -45 | 42 | 31 | 3.76 |
| Right inferior frontal gyrus | 48 | 12 | 30 | 24 | 3.97 |

**Table S32:** Encoding: High- vs. low-value cluster table

| *Region* | *x* | *y* | *z* | *Cluster size* | *z-max* |
|---|---|---|---|---|---|
| Superior parietal lobule / lateral occipital cortex / temporal occipital fusiform cortex / cerebellum | -33 | -54 | 51 | 4262 | 6.23 |
| Left frontal pole / inferior frontal gyrus / middle frontal gyrus | -51 | 42 | 9 | 1765 | 6.92 |
| Left caudate / thalamus | -18 | 12 | 6 | 232 | 5.67 |
| Right caudate | 18 | 18 | 12 | 54 | 4.17 |
| Right precentral gyrus | 51 | 9 | 33 | 50 | 4.26 |
| Right precentral gyrus | 24 | -9 | 54 | 47 | 4.79 |
| Left cerebellum | -3 | -51 | 0 | 38 | 4.73 |
| Right frontal pole | 51 | 39 | 12 | 35 | 4.53 |
| Right postcentral gyrus | 42 | -36 | 51 | 28 | 4.32 |
| Right putamen | 27 | 15 | -3 | 26 | 4.52 |
| Left thalamus | -3 | -24 | -3 | 23 | 4.65 |
| Left putamen | -30 | -15 | -6 | 20 | 5.7 |

**Table S33:** Encoding: High- vs. low-value by memory difference scores cluster table

| *Region* | *x* | *y* | *z* | *Cluster size* | *z-max* |
|---|---|---|---|---|---|
| Left lateral occipital cortex | -45 | -69 | -6 | 377 | 4.78 |
| Left middle frontal gyrus / inferior frontal gyrus | -48 | 21 | 27 | 232 | 4.95 |
| Right lateral occipital cortex (inferior) | 39 | -90 | 3 | 189 | 4.89 |
| Right temporal occipital fusiform cortex | 42 | -57 | -9 | 87 | 4.45 |
| Right lateral occipital cortex (superior) | 27 | -66 | 33 | 61 | 4.83 |

**Table S34:** Encoding: Remembered vs. not remembered cluster table

| ***Region*** | ***x*** | ***y*** | ***z*** | ***Cluster size*** | ***z-max*** |
|---|---|---|---|---|---|
| Right lateral occipital cortex / temporal occipital fusiform gyrus | 48 | -75 | -9 | 1296 | 5.75 |
| Left lateral occipital cortex / temporal occipital fusiform gyrus / inferior temporal gyrus | -48 | -72 | -6 | 1273 | 5.97 |
| Left inferior frontal gyrus | -48 | 9 | 27 | 103 | 4.22 |
| Left inferior frontal gyrus | -57 | 21 | -3 | 40 | 3.81 |
| Right hippocampus / amygdala | 24 | -6 | -21 | 21 | 4.06 |

**Table S35:** Retrieval: Retrieval vs. baseline by linear age cluster table

| ***Region*** | ***x*** | ***y*** | ***z*** | ***Cluster size*** | ***z-max*** |
|---|---|---|---|---|---|
| Right lateral occipital cortex | 45 | -81 | 21 | 147 | 5.28 |
| Left lateral occipital cortex | -39 | -87 | 21 | 59 | 4.25 |
| Right precentral gyrus / inferior frontal gyrus | 51 | 3 | 21 | 42 | 4.21 |
| Right precentral gyrus | 39 | -12 | 39 | 41 | 4.4 |
| Left postcentral gyrus / supramarginal gyrus | -63 | -24 | 48 | 38 | 4.45 |
| Left precentral gyrus | -60 | 6 | 21 | 36 | 4.39 |
| Right lateral occipital cortex | 27 | -60 | 57 | 34 | 3.76 |
| Cingulate gyrus / left thalamus | 3 | -33 | 0 | 30 | 4.74 |
| Left supramarginal gyrus | -69 | -24 | 24 | 28 | 4.33 |

**Table S36:** Retrieval: High- vs. low-value cluster table

| ***Region*** | ***x*** | ***y*** | ***z*** | ***Cluster size*** | ***z-max*** |
|---|---|---|---|---|---|
| Precuneus cortex | 0 | -72 | 39 | 299 | 4.81 |
| Left lateral occipital cortex | -33 | -69 | 51 | 236 | 5.28 |
| Left caudate / thalamus | -12 | -6 | 15 | 128 | 5.15 |
| Right cerebellum | 3 | -81 | -30 | 125 | 6.12 |
| Left inferior frontal gyrus / middle frontal gyrus | -48 | 21 | 24 | 116 | 4.57 |
| Left frontal orbital cortex | -30 | 30 | -3 | 88 | 4.82 |
| Right cerebellum | 39 | -69 | -36 | 85 | 4.65 |
| Right caudate | 15 | -3 | 21 | 77 | 4.96 |
| Left middle frontal gyrus | -45 | 12 | 48 | 74 | 4.31 |
| Cerebellum | -3 | -60 | -36 | 60 | 4.57 |
| Left inferior temporal gyrus | -57 | -60 | -15 | 60 | 4.33 |
| Cingulate gyrus | 0 | -33 | 3 | 25 | 4.09 |
| Right frontal orbital cortex | 33 | 33 | 3 | 21 | 4.69 |
| Left frontal pole | -36 | 57 | 3 | 20 | 3.79 |
| Right lateral occipital cortex | 27 | -66 | 42 | 20 | 4.24 |
